# How Functional Variants Reconfigure the Rac2 Conformational Landscape

**DOI:** 10.64898/2026.04.17.719286

**Authors:** Nurit Haspel, Hyunbum Jang, Ruth Nussinov

## Abstract

Rac2, a member of the Rho family of small GTPases, is a fundamental regulator of essential cellular processes. Pathogenic substitutions near and within the Switch II region, specifically D57N and E62K, have been implicated in oncogenesis and immunodeficiency. Despite their proximity, D57N is characterized as a loss-of-function mutation, while E62K is a constitutively active, gain-of-function mutation. In this study, we addressed several critical questions: (i) the structural basis of their altered cellular functions, (ii) how these variants rearrange the conformational ensemble, and (iii) the subsequent impact on cellular signaling networks. Using molecular dynamics (MD) simulations, we characterized the conformational dynamics of these Rac2 variants in GDP- and GTP-bound states. Our results demonstrate that Rac2^D57N^ predominantly adopts an inactive-like conformation, regardless of the bound nucleotide. GTP binding is insufficient to induce the canonical active state in this mutant. Conversely, Rac2^E62K^ maintains a nucleotide-dependent toggle, appearing inactive when bound to GDP and active when bound to GTP. Additionally, we examined the assembly of these variants with the regulator p50-RhoGAP. In the wild-type complex, GAP binding facilitates a shift toward a near-transition-state ensemble. In stark contrast, both the D57N and E62K complexes remain sequestered in a ground-ON state configuration, effectively trapping the GTPase and hindering GAP-mediated hydrolysis. While both Rac2 mutations result in immune system dysfunction, the underlying mechanisms are opposite: inactive vs. overactive. This work provides a high-resolution, mechanistic framework for understanding how localized perturbations in the switch loops landscape dictate systemic cellular outcomes.

## 1. Introduction

The Rho family, a subgroup of the Ras superfamily, regulates cellular activities such as cytoskeletal organization, gene expression, and transformation.^1–6^ Members of the Ras superfamily have been linked to multiple human cancers^6–13^ and act as molecular switches between active GTP-bound and inactive GDP-bound forms. The most well-characterized members of the Rho family are Rho, Rac, and Cdc42. The Rho protein has three isoforms: RhoA, RhoB, and RhoC. It controls the establishment of stress fibers, which generate the contractile force that propels the cell body behind the leading edge.^14^ Cdc42 is a key regulator of the actin cytoskeleton, controlling the formation of finger-like protrusions called filopodia.^15^ The three main mammalian Rac isoforms, Rac1, Rac2, and Rac3, have highly similar sequences and biochemical properties. Rac1 and Rac2 are very similar, sharing approximately 92% sequence identity, while Rac3 differs slightly. Rac1, in particular, controls the formation of lamellipodia, a crucial process for cell migration that enables cells to push their membranes forward.^16^ Rac2 is the focus of our current study. It plays a role in the biology of different cell types, such as neutrophils and lymphocytes,^17^ and was shown to be involved in several types of cancer and immunodeficiency disorders.^18–21^

Rac1 and Rac2 are composed of 192 amino acids (**Figure 1A**). Post-translational modifications include palmitoylation at Cys178 and geranylgeranylation at Cys189 for Rac1,^22^ while Rac2 is only modified by geranylgeranylation at Cys189. Rac2 consists of a catalytic domain (residues 1-177) and a hypervariable region (HVR, residues 178-192). The catalytic domain contains several highly conserved regions: the P-loop (residues 10-15), Switch I (resides 30-38), Switch II (residues 60-76), and the insert region (residues 122-135). The switch regions are responsible for interacting with GTPase activating proteins (GAPs) and guanine nucleotide exchange factors (GEFs)^23–25^ and undergo conformational changes upon guanosine activation and binding.^26^ GAPs negatively regulate GTPases by promoting the conversion of the active GTP-bound form to the inactive GDP-bound form.^27,28^ A highly conserved arginine finger (Arg282) of GAP protrudes into the GTP-binding site (**Figure 1B**) and interacts directly with GTP in the catalytic site of a GTPase.^29,30^ Mutation of this arginine residue of GAP reduces activity.^31^

**Fig. 1.**
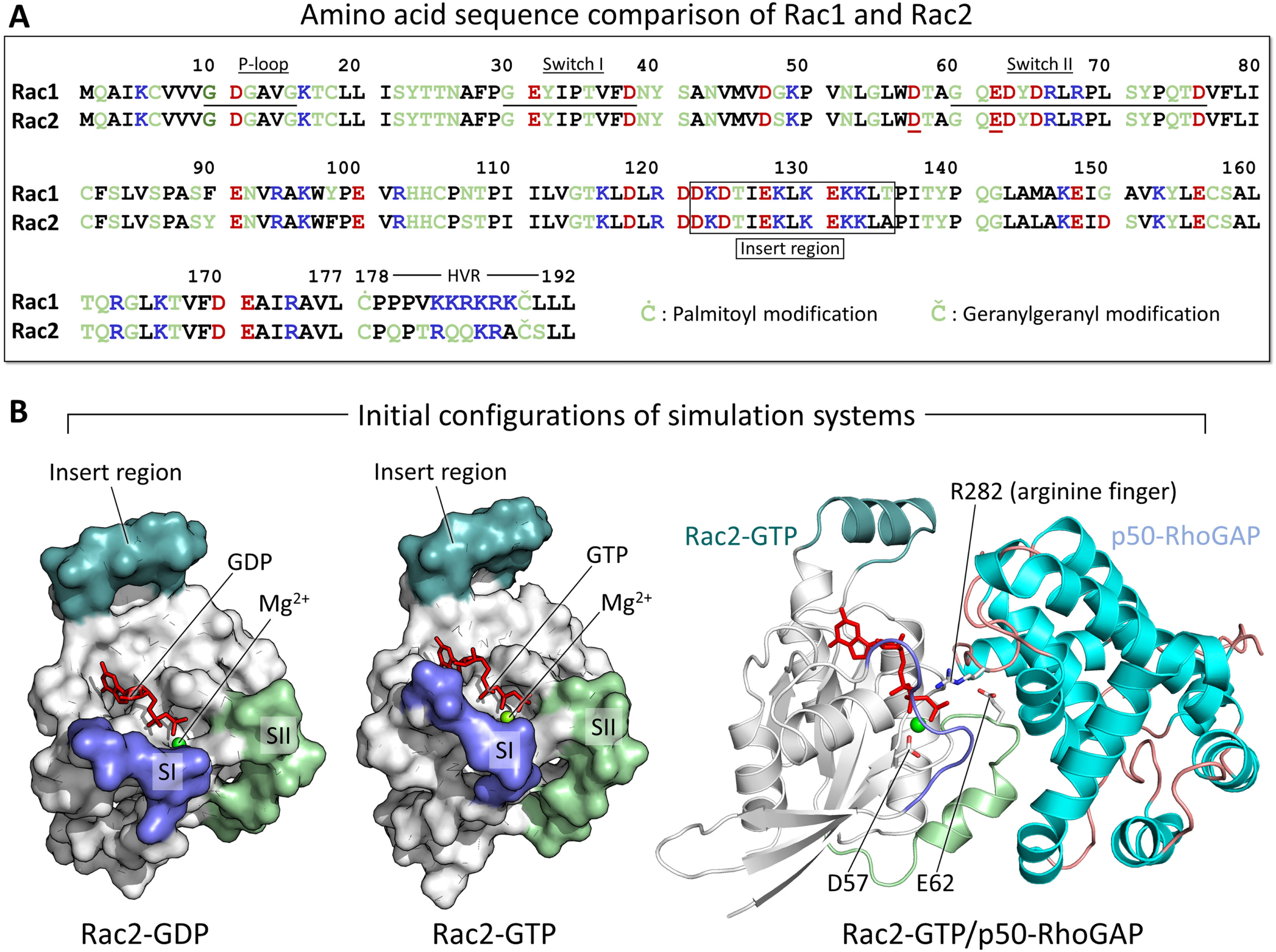
(A) Amino acid sequence comparison between Rac1 and Rac2 (*top panel*). In the sequence letter, hydrophobic, polar/glycine, positively charged, and negatively charged residues are colored black, green, blue, and red, respectively. (B) Initial configurations of isolated Rac2 in the GDP- and GTP-bound states, and Rac2-GTP in complex with p50-RhoGAP (*bottom panel*). SI and SII refer to Switch I and Switch II, respectively. In the complex, the mutation sites, Asp57 and Glu62, are marked in Rac2. The arginine finger, Arg282, is marked in p50-RhoGAP.

Several Rac2 mutations have been linked to cancer progression and immunodeficiency syndromes.^19,32,33^ G12R is associated with bone marrow hypoplasia.^34^ D57N is a dominant-negative mutation linked to neutrophil immunodeficiency syndrome^18,35,36^ and reduced superoxide production.^37^ E62K is an activating Rac2 mutation associated with lymphopenia, immunodeficiency, and cytoskeletal defects.^19,20^ Asp57 is located near the Switch II region, and Glu62 is located on it (**Figure 1B**). Asp57 is close to the nucleotide, and the mutation disrupts the binding with the nucleotide.^38^ Glu62 is at the binding interface of Rac2 with p50-RhoGAP (encoded by *ARHGAP1*). Therefore, both D57N and E62K affect in different ways despite their proximity. Both positions are highly conserved in the Ras and Rac families.^20^ These mutations are linked to cancer and autoimmunity through neutrophil dysfunction. The D57N mutation lies in a region that interacts directly with the GDP/GTP molecule. This results in a high rate of GTP dissociation which prevents activation,^19,33^ making it considered a loss-of-function mutation. E62K lies on the binding interface of Rac2 with p50-RhoGAP. The mutation disrupts the Rac2/GAP interaction and is therefore a gain-of-function mutation. It does not significantly change the Rac2 structure but causes Rac2 hyperactivation, alters GEF specificity (TIAM1 vs DOCK2), and impairs GAP function while retaining key effector interactions.^39^ The hyperactive Rac2^E62K^ variant was observed to induce cannibalistic cell-in-cell phenomena, specifically by promoting entosis in neighboring cells, thereby facilitating human immunodeficiency.^40,41^

To gain insight into the atomic-level differences between the wild type and the mutants, we performed all-atom molecular dynamics (MD) simulations to characterize the conformational ensembles and site-specific dynamics of Rac2 variants (D57N and E62K), in their monomeric forms and in complex with the p50-RhoGAP regulator. The D57N substitution destabilizes the salt bridge coordinating the Mg^2+^ cofactor for GTP. This results in a conformational shift that drives the Switch I and Switch II loops into an open conformation. With this open-switch loop architecture, GTP becomes increasingly exposed to the solvent. This structural instability correlates with the observed high rate of GTP dissociation, which is characteristic of a loss-of-function phenotype. Although association with p50-RhoGAP induces closure of the switch loops, the mutation’s intrinsic inability to stabilize negative charges at the active site prevents the complex from achieving a catalytically competent geometry. Consequently, the Rac2/GAP complex remains sequestered in a ground state, abrogating GAP-mediated hydrolysis. Conversely, the E62K variant favors a constitutively active state when bound to GTP, characterized by a stabilized closed switch loop configuration. Upon association with p50-RhoGAP, this closed state is maintained. However, charge inversion (Glu to Lys) disrupts the canonical interface between the Rac2 Switch II region and GAP, triggering significant rewiring of local salt bridge networks. This perturbs the conformational landscape, preventing the precise spatial coordination of catalytic atoms required for water-mediated GTP hydrolysis. Similar to the D57N variant, the E62K complex is trapped in a ground state, hindering GTP hydrolysis and leading to hyperactivation of downstream signaling. Unlike these pathogenic variants, the wild-type Rac2/GAP complex adheres to the established mechanical requirements for efficient GTP hydrolysis. The wild-type complex exhibits an inactive-like conformation of Rac2-GTP, indicating a near-transition-state ensemble. We present the mechanistic details of how the Rac2 variant reconfigures the conformational landscape and explain how these details are related to different clinical phenotypes.

## 2. Materials and Methods

### 2.1. Preparation of models

We simulated both isolated Rac2-GDP and Rac2-GTP, as well as Rac2-GTP in complex with p50-RhoGAP. For Rac2, we examined the D57N and E62K mutations and compared their conformations with those of the wild type. The Rac2-GDP coordinates were extracted from the crystal structure of Rac2-GDP/RhoGDI complex (PDB ID: 1DS6). The Rac2-GTP coordinates were adopted from the crystal structure of GTPγS-bound Rac2 with the G12V mutation (PDB ID: 2W2V). The GTPγS molecule was replaced with GTP, and the mutant residue was converted to the wild type. Then the wild-type Rac2-GTP was minimized to restore the P-loop conformation. The D57N and E62K mutant systems were constructed by replacing the residues Asp57 and Glu62 with Asn and Lys, respectively. Six isolated Rac2 systems were generated for the wild type (hereafter denoted as Rac2^WT^) and two mutants (hereafter denoted as Rac2^D57N^ and Rac2^E62K^) in the GDP- and GTP-bound states. Next, we formed a complex with p50-RhoGAP using Rac2^WT^-GTP and two mutants, Rac2^D57N^-GTP and Rac2^E62K^-GTP. However, since no crystal structure of the Rac2/p50-RhoGAP complex is available, we used the crystal structure of the RhoA-GDP/p50-RhoGAP complex in the transition state (PDB ID: 6R3V) to align Rac2-GTP with RhoA-GDP and replace it. Three systems, Rac2^WT^-GTP/p50-RhoGAP, Rac2^D57N^-GTP/p50-RhoGAP, and Rac2^E62K^-GTP/p50-RhoGAP complexes, were constructed for explicit MD simulations.

### 2.2. Simulation protocol

We followed the protocol outlined in previous studies.^28,42–47^ The simulations were conducted using the CHARMM^48^ all-atom additive force field (version C36m).^49,50^ The solvent was represented explicitly using the TIP3P model. The charge of all potential titratable groups was fixed at values corresponding to neutral pH (i.e. all acidic and basic side chains were represented in their negatively and positively charged forms, respectively). We used a cubic simulation box with periodic boundary conditions and the nearest image convention. The dimensions of the simulation box were set to 90 × 90 × 90 Å^3^ for the isolated Rac2 systems and 120 × 120 × 120 Å^3^ for the GAP-bound systems. Short-range van der Waals (vdW) interactions were calculated using switching function with twin-range cutoffs at 12 Å and 14 Å. To prevent discontinuities in the potential energy function, a smoothing factor was applied starting at a distance of 12.0 Å to ensure non-bonding energy terms smoothly converged to zero. Long-range electrostatic interactions were computed using the particle mesh Ewald (PME) method with a grid spacing of 1.0 Å. Na^+^ and Cl^−^ were added to neutralize each system and mimic physiological conditions. During the simulation, the potential energy was minimized by using 10,000 conjugate gradient steps. Then, the protein atoms were held fixed while the solvent was heated to a temperature of 310 K to ensure uniform distribution of the solvent around the protein. Next, the system was isothermally and isobarically equilibrated at 310 K and 1 bar (NPT conditions) to allow reaching infinite dilution conditions. Later, the solute was allowed to move, and the whole system was heated and equilibrated at the production temperature of 310 K and the pressure of 1 bar. The Langevin thermostat and the Nosé-Hoover Langevin piston control algorithm were employed to maintain constant temperature and pressure, respectively. The SHAKE algorithm was applied to constrain the motion of bonds involving hydrogen atoms. There are three replicas of each system, each of which was simulated for 1 µs. This results in an overall time of 27 µs. We used NAMD 2.14 for the production run simulations,^51^ and the results derived from these parallel trajectories showed consistent similarities, making them comparable. The integration time step was 2 fs for the production runs. For the isolated Rac2 systems, the root-mean-square-deviation (RMSD) revealed that Rac2 retained its initial structure, with RMSD values below 3 Å (**Figure S1**). Each replica simulation of Rac2/GAP complex yielded similar RMSD profiles for Rac2 and p50-RhoGAP.

## 3. Results

### 3.1. How Nucleotides and Mutations Reshape the Ensembles of Rac2 Conformation

We performed explicit MD simulations on the catalytic domains of wild-type Rac2 and its D57N and E62K mutants in both GDP- and GTP-bound states. During the simulations, we observed differences in the conformational flexibility of the Switch I and Switch II regions. To quantify these variations in their essential modes of motion, we performed an essential dynamics analysis on the six isolated Rac2 systems. Principal component analysis (PCA) reveals major divergence between the GDP- and GTP-bound systems, as well as between the mutant and wild-type systems (**Figure 2**). To further categorize these motions, we applied linkage clustering to the PCA projections and targeted two to three distinct clusters, as increasing the cluster count beyond three did not yield clusters of significant size. In the GDP-bound state, Rac2^WT^ forms three clusters, one of which is distinctly separated along PC1. In contrast, Rac2^D57N^ exhibits a single large cluster, while Rac2^E62K^ yields three overlapping clusters. In the GTP-bound state, both Rac2^WT^ and Rac2^E62K^ produce three overlapping clusters. However, Rac2^D57N^ displays two clusters clearly separated along PC1. The presence of non-overlapping clusters in these projections suggests discrete transitions between stable conformational states, highlighting the structural impact of the D57N and E62K mutations.

**Fig. 2.**
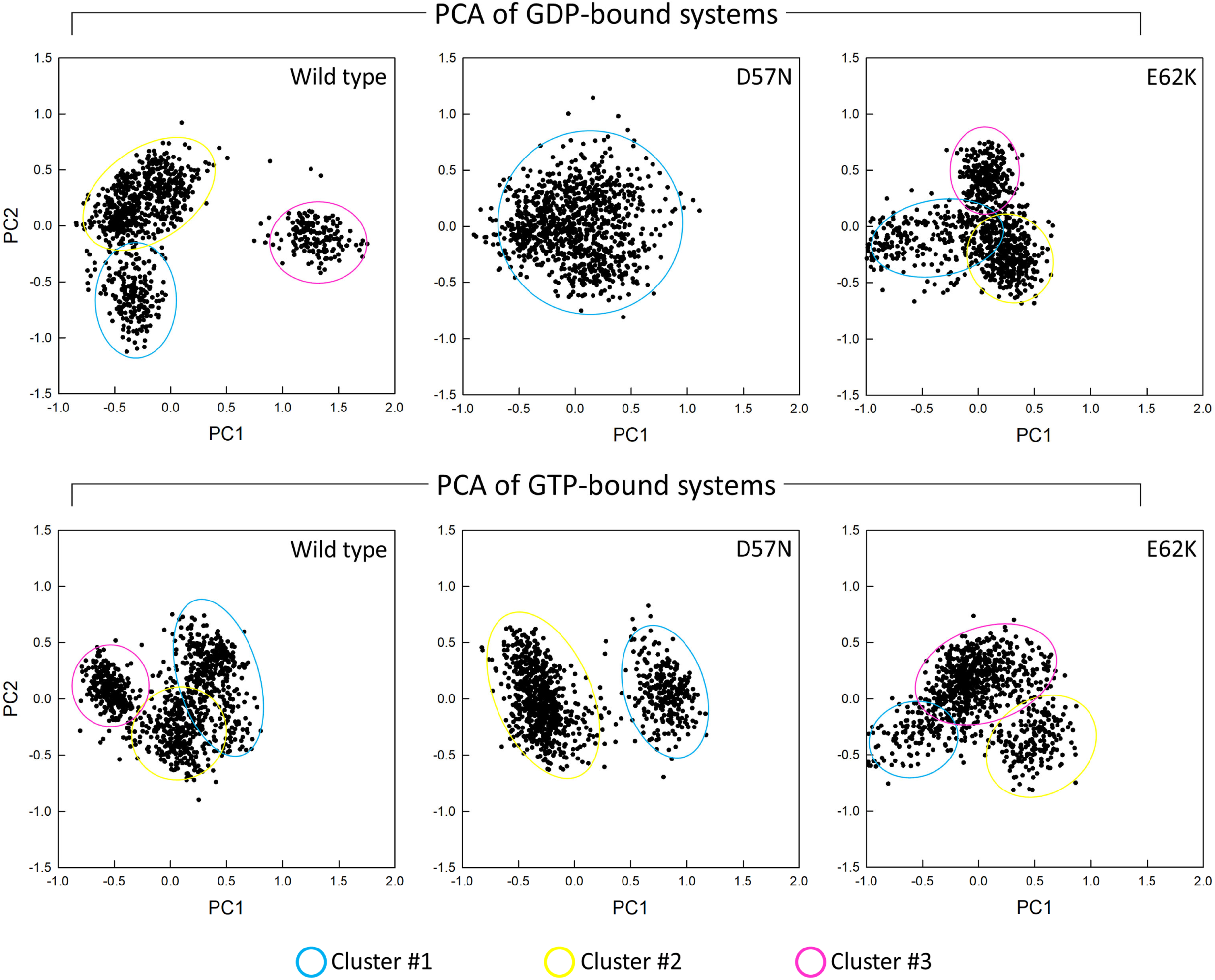
The projection of the first two principal components, PC1 and PC2 for the isolated Rac2^WT^, Rac2^D57N^, and Rac2^E62K^ in the GDP-bound (*top row*) and the GTP-bound (*bottom row*) states. Each cluster in the PCA projection was subject to linkage clustering and is shown in a different color.

To visualize the structural variations across clusters, we superimposed the representative conformations from each group (**Figure 3**). In all systems, the primary conformational divergences are localized to the Switch I and Switch II regions. In the GDP-bound systems, both Rac2^WT^ and Rac2^E62K^ exhibit distinct conformational shifts in the Switch I and Switch II loops between clusters; notably, the Switch II conformation in Rac2^WT^ Cluster 3 diverges significantly from the others. In contrast, Rac2^D57N^ exhibits no distinct conformational transitions in these loops. The average root-mean-square deviations (RMSDs) are comparable across the systems: 2.4 ± 0.4 Å for Rac2^WT^, 2.3 ± 0.2 Å for Rac2^D57N^, and 2.2 ± 0.2 Å for Rac2^E62K^. This suggests that loop reorientation drives these subtle variations in RMSD. In the GTP-bound systems, Rac2^WT^ displays only minor variations in the switch loops compared to the more pronounced changes in the D57N and E62K mutants. Average RMSDs, 2.1 ± 0.2 Å for Rac2^WT^, 2.3 ± 0.2 Å for Rac2^D57N^, and 2.4 ± 0.3 Å for Rac2^E62K^, remain similar to their GDP-bound counterparts. While the overall structural fold is preserved, high dynamic fluctuations are concentrated within the switch regions. To quantify these local dynamics, we calculated the root-mean-square fluctuations (RMSFs) per residue (**Figure S2**). Both switch regions exhibit higher flexibility than the protein core across all systems. In the wild-type and Rac2^D57N^-GTP systems, Switch II generally fluctuates more than Switch I. However, Rac2^D57N^-GDP and Rac2^E62K^-GDP/GTP exhibit slightly suppressed fluctuations. Notably, the RMSFs of the insert regions in all Rac2 systems are lower than those reported for Cdc42,^47^ suggesting that these proteins undergo distinct conformational dynamics despite belonging to the same Rho GTPase family.

**Fig. 3.**
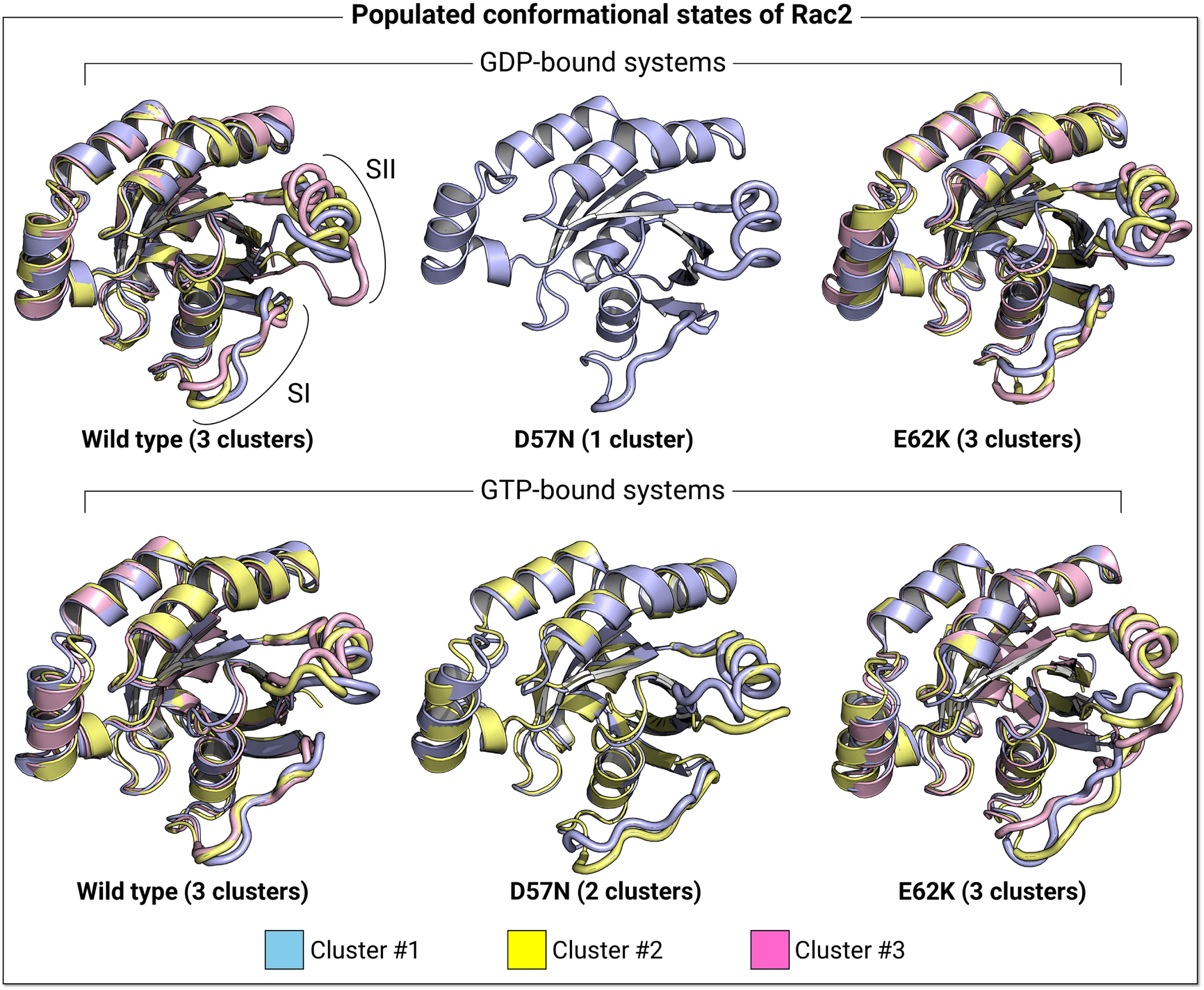
Snapshots representing the superimposition of the best representative conformations from the clustering for the isolated Rac2^WT^, Rac2^D57N^, and Rac2^E62K^ in the GDP-bound (*top row*) and the GTP-bound (*bottom row*) states. The protein formations are depicted from the linkage clustering in the PCA projection. Both switch regions are shown as thick tubes. SI and SII refer to Switch I and Switch II, respectively.

To map the conformational landscape of the switch loops, we calculated the probability distributions for two defining atom-pair distances: *d*_1_ (defined by the distance from the Cα atom of Gly60 in Switch II to the β-phosphate (P_β_) of GDP/GTP) and *d*_2_ (defined by the distance from the Cα atom of Thr35 in Switch I to the P_β_ of GDP/GTP). The resulting two-dimensional potential of mean force (PMF) contour plot reveals that Rac2^WT^-GDP prefers an open Switch II configuration, whereas the GTP-bound form shifts toward a closed state (**Figure 4**). Normal mode analysis (NMA) corroborates these findings; the lowest-frequency motions show large displacements in the Switch II loop of the inactive GDP-bound form, while the active GTP-bound state exhibits suppressed fluctuations (**Figure S3**). In contrast, Rac2^D57N^ adopts an open Switch I configuration with high fluctuations in both GDP- and GTP-bound states. Notably, the GTP-bound form displays an even more pronounced Switch I opening than the GDP-bound state, resulting in a conformation characteristic of an inactive state. In the GDP-bound state, Rac2^E62K^ adopts an open Switch I configuration, indicative of an inactive state. Conversely, the GTP-bound form promotes the closure of both Switch I and II loops, yielding an active conformation. It is important to note that for the D57N mutation, GTP binding does not necessarily produce an active conformation. This atypical behavior mirrors the conformational transitions observed in the K-Ras4B G12V mutant when bound to GDP.^52^

**Fig. 4.**
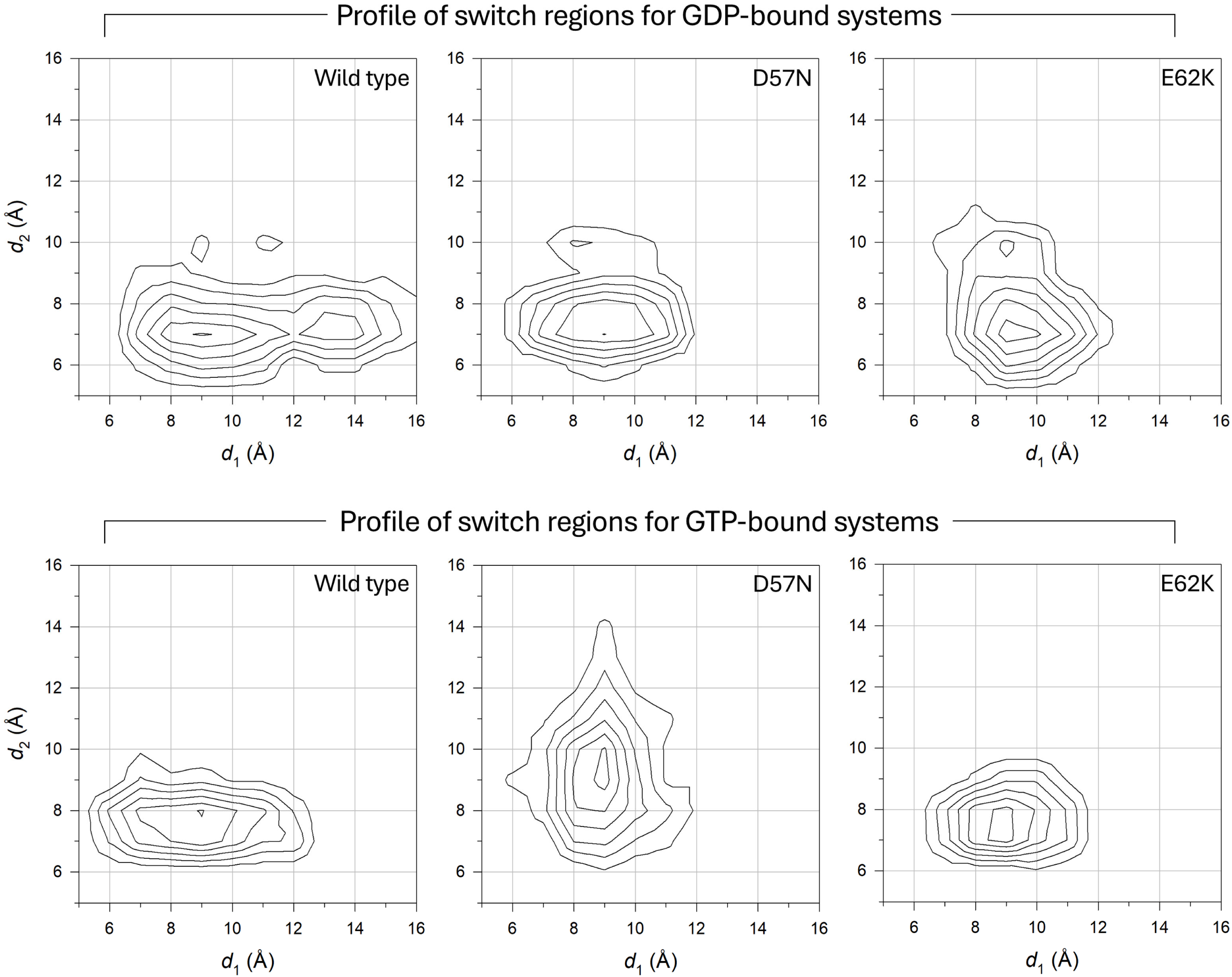
The two-dimensional potential of mean force, Δ*G*_PMF_, representing the relative free energy profile along reaction coordinates, *d*_1_ and *d*_2_. In the calculation of the probability distributions for two atom pair distances, *d*_1_ is defined as the distance from Gly60-Cα to GTP/GDP-P_β_ and *d*_2_ is defined as the distance from Thr35-Cα to GTP/GDP-P_β_. The contour plots illustrate the distance profiles for isolated Rac2^WT^, Rac2^D57N^, and Rac2^E62K^ in the GDP-bound (*top row*) and the GTP-bound (*bottom row*) states. The contour plot of Rac2^D57N^-GTP is highly analogous to that of its GDP-bound form, which suggests an inactive state.

### 3.2. The D57N Mutation Alters the Conformational Landscape of GTP-bound Rac2, Favoring an Inactive-like Ensemble

To elucidate the structural basis of the inactive-like conformation of Rac2^D57N^-GTP, we analyzed the interatomic distances between the Asp57 (or Asn57 for mutant) residue and Mg^2+^ in the GTP-bound state. In the wild-type active site, Mg^2+^ is coordinated through electrostatic interactions with Thr17 and Asp57, and GTP is stabilized via a salt bridge with Lys16 (**Figure 5**). Comparative analysis of all systems reveals that the Thr17–Mg^2+^ coordination and the Lys16–GTP salt bridge are highly conserved (**Figure S4**). However, the Asn57–Mg^2+^ distance in the Rac2^D57N^ mutant is increased relative to the corresponding distances in both Rac2^WT^ and Rac2^E62K^. The average Asp57–Mg^2+^ distances are 4.5 ± 0.2 Å and 4.6 ± 0.2 Å in Rac2^WT^ and Rac2^E62K^, respectively. The corresponding average Asn57–Mg^2+^ distance in Rac2^D57N^ is 5.3 ± 0.3 Å. Substituting Asp with Asn disrupts the critical coordination of Mg^2+^, thereby destabilizing the catalytic pocket. This loss of structural constraint triggers a conformational shift, resulting in the characteristic open conformation of the Switch I loop. The transitional behavior of Rac2^D57N^-GTP indicates an instability of the active site that remains in an inactive-like state. This structural instability confirms that the D57N mutation acts as a dominant-negative variant, resulting in a loss of function.

**Fig. 5.**
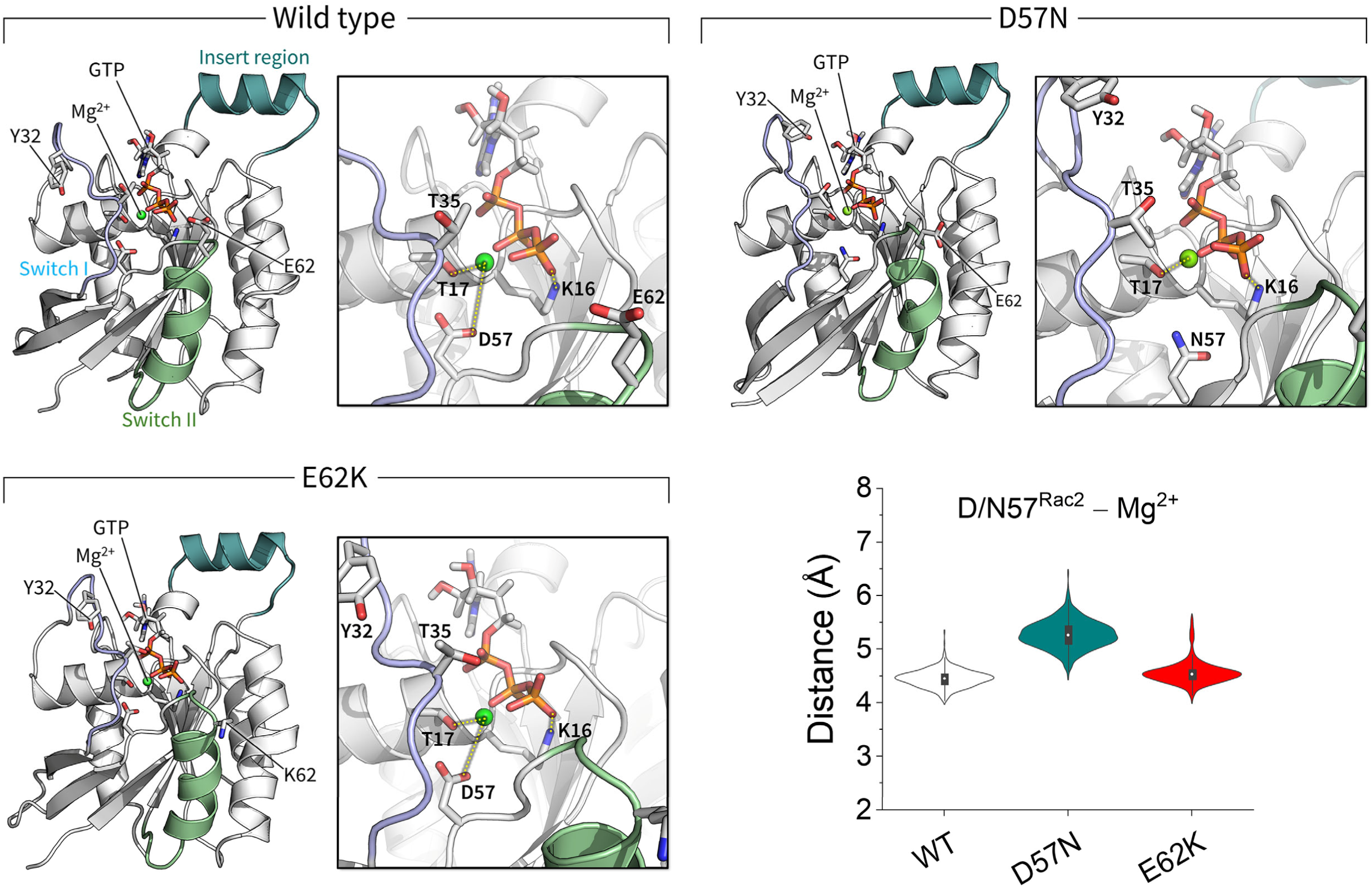
Snapshots highlighting the coordination of Thr17 and Asp/Asn57 with Mg^2+^ and of Lys16 with the Pγ of GTP for the isolated Rac2^WT^, Rac2^D57N^, and Rac2^E62K^ in the GTP-bound state. The violin plot shows the atomic pair distance between Asp57-CG (Asn57-CG for the D57N mutant) and Mg^2+^ (*bottom right*). The mutant residue Asn57 impairs the salt bridge with Mg^2+^, leading to the destabilization of GTP at the active site.

### 3.3. D57N and E62K Mutations Abrogate Rac2 Sensitivity to GAP Activity

Small GTPases are inactivated by GAPs, which catalyze the hydrolysis of GTP to GDP. By accelerating the slow intrinsic rate of hydrolysis, GAPs serve as essential negative regulators of downstream signaling cascades. To elucidate the biochemical implications of specific Rac2 variants, we performed all-atom explicit-solvent MD simulations on three distinct complexes: Rac2^WT^-GTP/p50-RhoGAP, Rac2^D57N^-GTP/p50-RhoGAP, and Rac2^E62K^-GTP/p50-RhoGAP. The initial p50-RhoGAP coordinates were standardized across all three systems to enable direct comparison of the D57N and E62K mutations. Although p50-RhoGAP exhibited localized fluctuations at the Rac2 interface during the simulations, the complex remained stable with no dissociation events observed (**Figure S5**). The stability of the complex conformation is maintained by an intermolecular salt bridge between Asp63^Rac2^ and Lys319^GAP^, which is highly conserved across all systems (**Figure 6**). Both the wild-type and D57N systems exhibit similar salt bridge profiles at the interface. During the simulations, the salt bridges, Glu62^Rac2^–Arg323^GAP^ and Asp63^Rac2^–Arg323^GAP^ are sustained, while the Asp65^Rac2^–Arg323^GAP^ interaction is diminished. In contrast, the E62K mutation disrupts the interfacial salt bridge network. The charge-reversal mutation to Lys62^Rac2^ eliminates electrostatic attractions with Arg291^GAP^, Lys319^GAP^, and Arg323^GAP^. Instead, a new salt bridge is established between Lys62^Rac2^ and Glu324^GAP^. While the Asp65^Rac2^–Arg323^GAP^ interaction is sustained, the Asp63^Rac2^–Arg323^GAP^ interaction is diminished during the simulation. Similar salt bridge interactions involving the Switch II residues, Glu62 and Glu63, were also identified in the K-Ras4B/NF1 complex.^28^ While the Switch I residues of K-Ras4B are primarily responsible for forming salt bridges with NF1, in Rac2, only the Switch I residue Glu31 weakly forms a salt bridge with p50-RhoGAP. Instead, the insert region residue Lys132 forms a weak salt bridge, which cannot be observed in K-Ras4B. Taken together, the distinct primary sequences and topologies of these GAPs (**Figure S6**) suggest a high level of substrate specificity, implying that p50-RhoGAP may be unable to activate Ras, and *vice versa*.

**Fig. 6.**
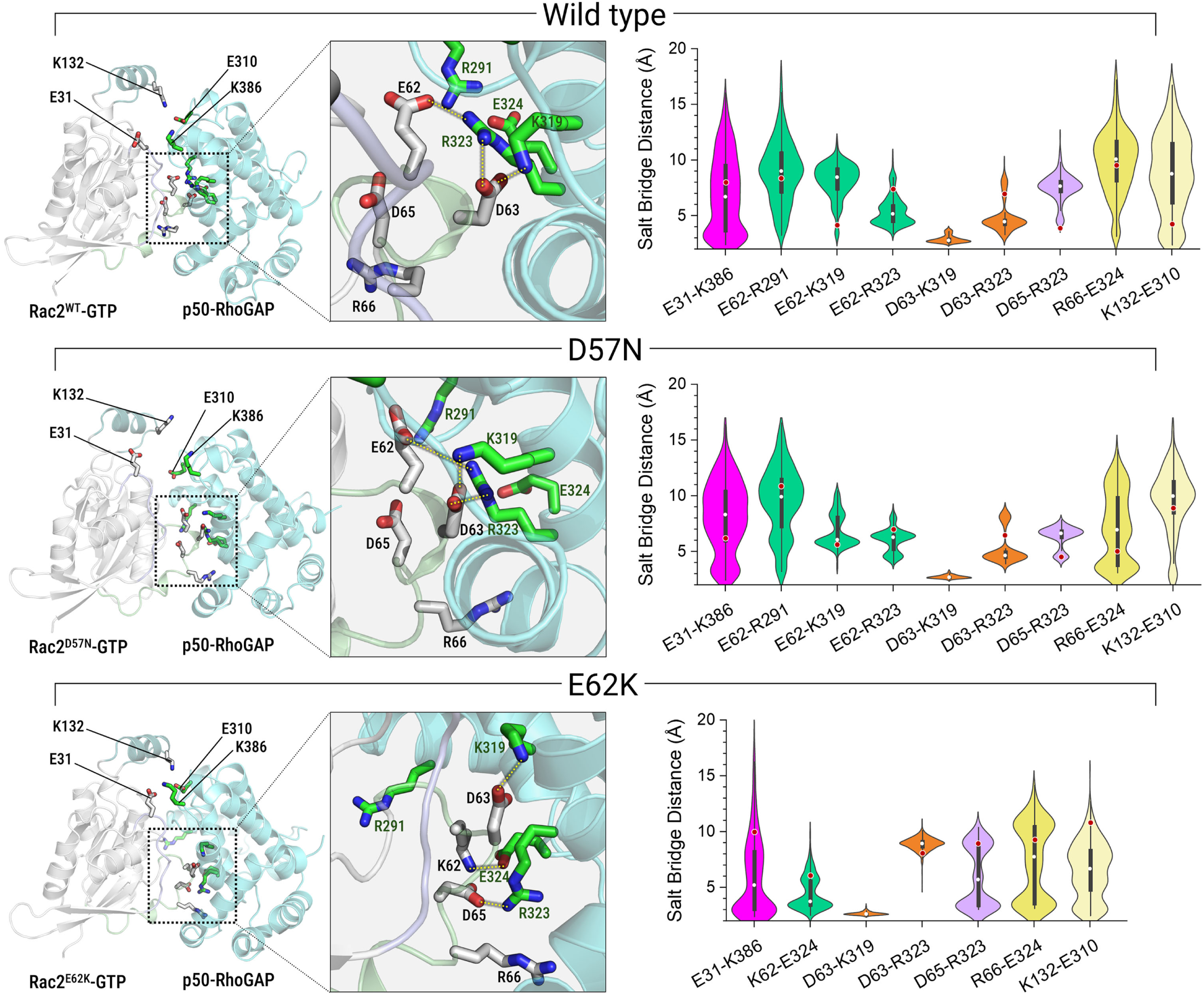
Comparative snapshots of interfacial salt bridge formations between Rac2 and p50-RhoGAP for the GTP-bound wild-type Rac2 and its D57N and E62K mutants in complex with p50-RhoGAP. The highlighted interactions focus on the Switch II region at the protein-protein interface. The violin plots show the salt bridge distances between potential atomic pairs at the interface. The mutant residue Lys62^Rac2^ disrupts the salt bridge network, rewiring its interaction with Glu324^GAP^ instead.

GTP hydrolysis occurs when a GTP-bound GTPase forms a complex with its specific GAP. This catalytic event involves the transition of GTPase/GAP complex from the ground state to the transition state, which is defined by the movement of a conserved arginine finger from the GAP. In the ground-OFF state, the arginine finger is oriented away from the active site of the GTPase. Then, in the ground-ON state, it reorients into the active site to coordinate with the γ-phosphate (Pγ) of the GTP molecule. While GTPases maintain an active conformation during the ground states, the transition state subsequently adopts a loose, dissociative-like geometry. This geometry facilitates cleavage of the β-γ phosphoanhydride bond and subsequent proton transfer from a water molecule to the Pγ of GTP.^53,54^ In our simulations, the wild-type Rac2^WT^-GTP/p50-RhoGAP complex displayed a catalytic environment consistent with established GTP hydrolysis mechanisms (**Figure 7**). This near-transition-state ensemble is characterized by the deep insertion of the arginine finger (Arg282) into the active site, stable coordination of Mg^2+^ by both Thr17 and Asp57, formation of a salt bridge between Lys16 and Pγ, and coordination of Gln61 at the active site, which facilitates the accumulation of residual water molecules at the cleavage site. In contrast, Rac2 mutations disrupt these canonical interactions. In the Rac2^D57N^-GTP/p50-RhoGAP complex, the mutant residue Asn57 causes a total loss of Mg^2+^ coordination. The average Asn57–Mg^2+^ distance is 5.4 ± 0.5 Å, which is similar to that of the isolated Rac2^D57N^ system. This results in the shallow insertion of the Arg282 side chain into the active site due to an electrostatic shift, likely reducing the stabilization of the transition state. Conversely, the corresponding average Asp57–Mg^2+^ distances are 3.0 ± 0.1 Å and 2.3 ± 0.1 Å in Rac2^WT^ and Rac2^E62K^, respectively, in complex with p50-RhoGAP. These distances are much shorter than those observed in their isolated systems. In the Rac2^E62K^-GTP/p50-RhoGAP complex, the E62K substitution induces a conformational shift that disrupts the canonical catalytic architecture. Thr17 loses coordination with Mg^2+^, and Gln61 changes the orientation of its side chain away from Pγ. These changes suggest that the mutant residue Lys62 shifts the GAP pose with respect to Rac2 due to a weakened Switch II interaction, thereby rewiring the salt bridge interactions between Switch II and GAP.

**Fig. 7.**
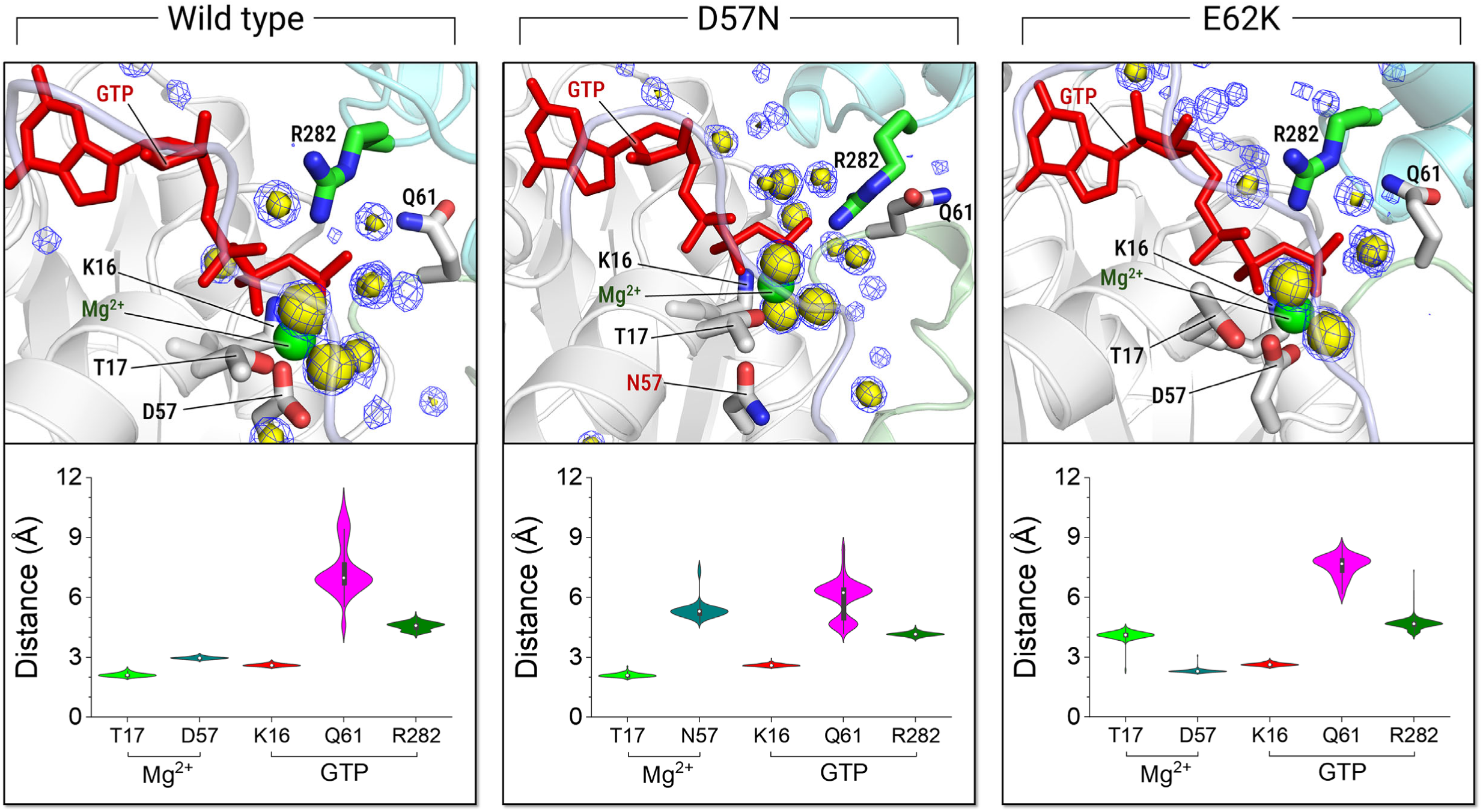
Coordination geometry and hydration of the Rac2/p50-RhoGAP active site. Snapshots highlighting the coordination of Thr17 and Asp/Asn57 with Mg^2+^ and the coordination of Lys16, Gln61, and the Arg282 finger with the Pγ of GTP in the wild-type Rac2 and its D57N and E62K mutants in complex with p50-RhoGAP. In the snapshots, water molecules are represented by a three-dimensional density map that reveals hydration in the active site, with probabilities *P* = 0.6 (yellow surface) and *P* = 0.5 (blue mesh). The violin plots show the atomic pair distances of Thr17-OG1 and Asp57-CG (Asn57-CG for the D57N mutant) from Mg^2+^ and of Lys16-NZ, Gln61-CD, and Arg282-CZ from the Pγ of GTP.

The initial architecture of the complex was modeled by adopting the transition-state GAP conformation from the crystal structure (PDB ID: 6R3V) and aligning the active, GTP-bound Rac2 onto it. The resulting assembly initially exhibited a ground-ON state configuration, characterized by an “IN" orientation of the Arg282 finger and a closed Switch I conformation. In the wild-type complex, while the arginine finger remained stably inserted in the active site, Rac2 exhibited increased flexibility in the Switch I loop during the simulations, resulting in an open conformation (**Figure S7**). This shift suggests that the wild-type complex effectively samples ensembles corresponding to the transition-like state, which facilitates the catalytic pathway. In stark contrast, both the D57N and E62K mutations appear to restrict these essential dynamics. In these mutant complexes, the GAP induces or stabilizes a closed Switch I loop conformation compared to the isolated systems (**Figure 4**). This suggests that the mutations trap the complexes in a catalytically unproductive ground-ON state, which may hinder GAP-mediated GTP hydrolysis. While MD simulations cannot directly capture this catalytic process—requiring kinetic studies or QM simulations for explicit modeling—this mechanism is strongly supported by the observed mutation-induced conformational changes and the loss of key pairwise interactions.

## 4. Discussion

We employed MD simulations to characterize the structural and dynamical landscapes of Rac2 with the D57N and E62K mutations in both GDP- and GTP-bound states. Our objective was to elucidate how these mutations perturb the conformational ensembles governing the transition between active and inactive states.^55–57^ The primary structural divergence between the inactive (GDP-bound) and active (GTP-bound) wild-type Rac2 resides in the Switch I and Switch II loops. In the inactive state, these loops adopt an open configuration relative to the nucleotide-binding pocket, whereas GTP binding induces a closed conformation that stabilizes the active site. Our simulations reveal that the D57N and E62K mutations alter these transitions through distinct switch loop dynamics. The D57N substitution, associated with a loss-of-function phenotype, replaces an Asp with a neutral Asn. This alteration impairs the essential coordination of the Mg^2+^ cofactor for GTP, leading to a destabilized active site. Even in the presence of GTP, the D57N variant favors an ensemble resembling the GDP-bound Rac2, characterized by open Switch I and Switch II loops. This conformational shift toward an inactive state likely underlies the dominant-negative effect by preventing productive interactions with effectors. Previous studies^58^ have shown that the D57A mutation in N-Ras-GTP, which is related to melanoma, has a similar effect on the active site, disrupting the interaction with Mg^2+^. Conversely, the E62K mutation in Rac2-GTP functions through a gain-of-function mechanism. The inversion of charge, from the negatively charged Glu to the positively charged Lys, appears to enhance the stability of the active-site architecture. This electrostatic shift stabilizes the complex, promoting a persistently closed conformation of both switch loops. Consistent with its observed hyper-activating clinical profile, Rac2^E62K^-GTP prolongs its active state.

GAPs function as critical negative regulators of GTPases by accelerating the hydrolysis of GTP to GDP, effectively transitioning the protein from an active to an inactive state. Binding of the effector (or regulator) molecule promotes an allosteric conformational change, shifting the conformational ensemble from a dormant (inactive) state to an active state, or vice versa.^55–57,59,60^ Rac2, a Rho-family GTPase, is essential for the functional integrity of neutrophils and lymphocytes. Pathogenic mutations in Rac2 are frequently implicated in oncogenesis and primary immunodeficiency disorders. Notably, the D57N mutation, located within the Switch II loop, is a dominant-negative variant responsible for neutrophil immunodeficiency syndrome.^17–19^ The E62K mutation, also located within the Switch II loop, is involved in the interaction at the Rac2/GAP binding interface, clinically associated with lymphopenia, immunodeficiency, and profound cytoskeletal defects.^20,40^ Through detailed structural investigations, our study elucidates the mechanisms by which pathogenic Rac2 variants allosterically alter both conformational dynamics and GAP-mediated regulation. Specifically, our analysis suggests that Rac2^D57N^ and Rac2^E62K^ variants disrupt GAP-mediated GTP hydrolysis, thereby impairing the intrinsic suppressing activity of p50-RhoGAP towards the Rac2 GTPase.

The association of Rac2^WT^-GTP with p50-RhoGAP shifts the conformational ensemble toward a GDP-bound-like inactive state, which lowers the energetic barrier for GTP hydrolysis. This catalytic priming is driven by the precise spatial organization of the catalytic groups at the active site. The highly conserved Arg282 finger of the GAP inserts into the Rac2 active site and coordinates with the Pγ of GTP. This interaction stabilizes the negative charges on the β-γ bridging oxygen of GTP. GAP-binding optimally positions Gln61, which subsequently recruits catalytic water molecules to coordinate with the Pγ. This highly coordinated atomic architecture induces a global conformational rearrangement within Rac2, characterized by a transition toward open Switch I and Switch II loops, similar to the inactive state. Our MD observations suggest that the Rac2^WT^-GTP/p50-RhoGAP complex effectively samples the transition-state ensemble, rendering it highly susceptible to GTP hydrolysis. Notably, while classical MD cannot directly simulate the nucleophilic attack and subsequent hydrolysis, it provides the key geometric criteria that strongly imply it.

In the Rac2^D57N^-GTP/p50-RhoGAP complex, GAP recruitment induces closure of the Rac2 switch loops, a marked contrast from the open configuration observed in the Rac2^D57N^-GTP monomer. However, despite the proper orientation of the Arg282 arginine finger of GAP within the active site, the mutation’s intrinsic inability to stabilize the localized negative charge density precludes the formation of a catalytically competent transition state. The Rac2^E62K^-GTP/p50-RhoGAP complex maintains the stabilized closed switch loop configuration seen in its monomeric form. However, the mutant residue Lys62 disrupts the canonical salt bridge networks at the interface between Switch II and GAP. This rearranged conformational landscape prevents the precise spatial coordination of the catalytic atoms at the active site and water required for the catalytic event. Both the D57N and E62K variants effectively abrogate GAP-mediated hydrolysis, hinder GTP hydrolysis by GAP, retaining the protein in the GTP-bound state. Despite the “IN” position of the Arg282 finger, both mutant complexes remain trapped in a ground-ON state configuration. This creates a formidable intrinsic energy barrier that prevents the system from reaching the transition state necessary for catalysis.

Although both variants result in an accumulation of GTP-bound Rac2, they drive radically opposed clinical phenotypes through distinct molecular mechanisms. For the loss-of-function Rac2^D57N^ variant, its primary defect is an impaired affinity for GTP and a high dissociation rate. This renders the protein signaling-incompetent and allows it to interfere with wild-type Rac2 function, leading to severe neutrophil dysfunction. For the gain-of-function Rac2^E62K^ variant, it is hyperactivated. It binds GTP stably but cannot hydrolyze it, leading to constitutive signaling causing immune dysregulation. By resolving how these variants reconfigure the conformational landscape, we provide a high-resolution structural framework that explains how localized amino acid substitutions translate into these contrasting clinical syndromes. These results provide a mechanistic benchmark, illuminating how specific pathogenic mutations disrupt the intricate interplay between Rac2, the bound nucleotide, and the GAP to alter enzymatic activity.

Although efforts are underway to design Rac2 specific drugs for cancer and immune diseases there are currently no specific FDA-approved drugs. We anticipate that our findings, particularly regarding conformational changes in the switch regions, the critical roles of specific amino acids in Rac2-GTP binding, and the effect of point mutations on Rac2/GAP interactions, will inspire future drug discovery efforts.

## Author Contributions

N.H., H.J, and R.N. conceived and designed the study. N.H. and H.J performed MD simulations and prepared the first draft. N.H. and H.J. analyzed the data, generated figures, and wrote the manuscript. All authors edited and approved the manuscript.

## Notes

The authors declare no competing financial interest.

## Supporting Information

Figures for RMSDs of three replica simulations for all systems; RMSFs of isolated Rac2 in the GDP- and GTP-bound states; stereo views of superimpositions of protein motion at the main normal mode with the lowest-frequency motion; salt bridge distances representing atom coordination at the active site; snapshots of Rac2 in complex with superimpositions of p50-RhoGAP; comparisons of the sequences and structures of NF1 GRD and p50-RhoGAP; contour plots representing the distance profiles of the switch loop with respect to the nucleotides.

## Acknowledgments

This Research was supported by the Cancer Innovation Laboratory, Center for Cancer Research, National Cancer Institute, National Institutes of Health Intramural Research Program project number ZIA BC 010441 and federal funds from the National Cancer Institute, National Institutes of Health, under contract HHSN261201500003I. The contributions of the NIH authors were made as part of their official duties as NIH federal employees, are in compliance with agency policy requirements, and are considered Works of the United States Government. However, the findings and conclusions presented in this paper are those of the authors and do not necessarily reflect the views of the NIH or the U.S. Department of Health and Human Services. The simulations have been performed using the high-performance computational facilities of the Biowulf PC/Linux cluster at the National Institutes of Health, Bethesda, MD (https://hpc.nih.gov/), on the UMass Boston research cluster and the Unity research cluster at the Massachusetts High Performance Computing Center (MGHPCC) (https://unityhpc.org/).

## TOC Graphic

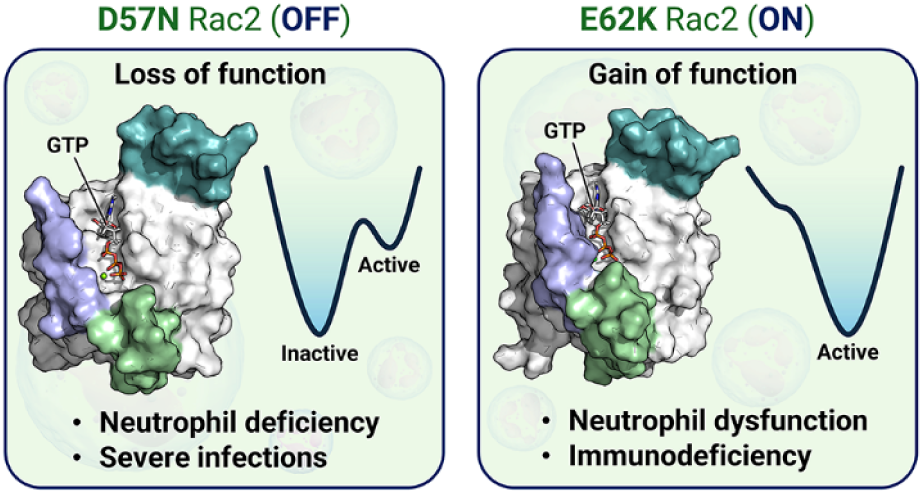

## References

(1) Cardama, G. A.; Gonzalez, N.; Maggio, J.; Menna, P. L.; Gomez, D. E. Rho GTPases as therapeutic targets in cancer (Review). Int J Oncol 2017, 51, 1025–1034. DOI: 10.3892/ijo.2017.4093.

(2) Ridley, A. J. Rho GTPases and cell migration. J Cell Sci 2001, 114, 2713–2722. DOI: 10.1242/jcs.114.15.2713.

(3) Owen, D.; Campbell, L. J.; Littlefield, K.; Evetts, K. A.; Li, Z.; Sacks, D. B.; Lowe, P. N.; Mott, H. R. The IQGAP1-Rac1 and IQGAP1-Cdc42 interactions: interfaces differ between the complexes. J Biol Chem 2008, 283, 1692–1704. DOI: 10.1074/jbc.M707257200.

(4) Jang, H.; Muratcioglu, S.; Gursoy, A.; Keskin, O.; Nussinov, R. Membrane-associated Ras dimers are isoform-specific: K-Ras dimers differ from H-Ras dimers. Biochem J 2016, 473, 1719–1732. DOI: 10.1042/BCJ20160031.

(5) Narumiya, S.; Thumkeo, D. Rho signaling research: history, current status and future directions. FEBS Lett 2018, 592, 1763–1776. DOI: 10.1002/1873-3468.13087.

(6) Liu, L.; Zhang, L.; Zhao, S.; Zhao, X. Y.; Min, P. X.; Ma, Y. D.; Wang, Y. Y.; Chen, Y.; Tang, S. J.; Zhang, Y. J.; Du, J.; Gu, L. Non-canonical Notch Signaling Regulates Actin Remodeling in Cell Migration by Activating PI3K/AKT/Cdc42 Pathway. Front Pharmacol 2019, 10, 370. DOI: 10.3389/fphar.2019.00370.

(7) Nussinov, R.; Jang, H. Direct K-Ras Inhibitors to Treat Cancers: Progress, New Insights, and Approaches to Treat Resistance. Annu Rev Pharmacol Toxicol 2024, 64, 231–253. DOI: 10.1146/annurev-pharmtox-022823-113946.

(8) Bustelo, X. R. RHO GTPases in cancer: known facts, open questions, and therapeutic challenges. Biochem Soc Trans 2018, 46, 741–760. DOI: 10.1042/BST20170531.

(9) Maldonado, M. D. M.; Dharmawardhane, S. Targeting Rac and Cdc42 GTPases in Cancer. Cancer Res 2018, 78, 3101–3111. DOI: 10.1158/0008-5472.CAN-18-0619.

(10) Maldonado, M. D. M.; Medina, J. I.; Velazquez, L.; Dharmawardhane, S. Targeting Rac and Cdc42 GEFs in Metastatic Cancer. Front Cell Dev Biol 2020, 8, 201. DOI: 10.3389/fcell.2020.00201.

(11) Crosas-Molist, E.; Samain, R.; Kohlhammer, L.; Orgaz, J. L.; George, S. L.; Maiques, O.; Barcelo, J.; Sanz-Moreno, V. Rho GTPase signaling in cancer progression and dissemination. Physiol Rev 2022, 102, 455–510. DOI: 10.1152/physrev.00045.2020.

(12) Clayton, N. S.; Ridley, A. J. Targeting Rho GTPase Signaling Networks in Cancer. Front Cell Dev Biol 2020, 8, 222. DOI: 10.3389/fcell.2020.00222.

(13) Haga, R. B.; Ridley, A. J. Rho GTPases: Regulation and roles in cancer cell biology. Small GTPases 2016, 7, 207–221. DOI: 10.1080/21541248.2016.1232583.

(14) Hall, A. Rho GTPases and the actin cytoskeleton. Science 1998, 279, 509–514. DOI: 10.1126/science.279.5350.509.

(15) Ahmed, S.; Goh, W. I.; Bu, W. I-BAR domains, IRSp53 and filopodium formation. Semin Cell Dev Biol 2010, 21, 350–356. DOI: 10.1016/j.semcdb.2009.11.008.

(16) Ma, N.; Xu, E.; Luo, Q.; Song, G. Rac1: A Regulator of Cell Migration and a Potential Target for Cancer Therapy. Molecules 2023, 28, 2976. DOI: 10.3390/molecules28072976.

(17) Lougaris, V.; Baronio, M.; Gazzurelli, L.; Benvenuto, A.; Plebani, A. RAC2 and primary human immune deficiencies. J Leukoc Biol 2020, 108, 687–696. DOI: 10.1002/JLB.5MR0520-194RR.

(18) Ambruso, D. R.; Knall, C.; Abell, A. N.; Panepinto, J.; Kurkchubasche, A.; Thurman, G.; Gonzalez-Aller, C.; Hiester, A.; deBoer, M.; Harbeck, R. J.; Oyer, R.; Johnson, G. L.; Roos, D. Human neutrophil immunodeficiency syndrome is associated with an inhibitory Rac2 mutation. Proc Natl Acad Sci U S A 2000, 97, 4654–4659. DOI: 10.1073/pnas.080074897.

(19) Hsu, A. P. Not too little, not too much: the impact of mutation types in Wiskott-Aldrich syndrome and RAC2 patients. Clin Exp Immunol 2023, 212, 137–146. DOI: 10.1093/cei/uxad001.

(20) Hsu, A. P.; Donko, A.; Arrington, M. E.; Swamydas, M.; Fink, D.; Das, A.; Escobedo, O.; Bonagura, V.; Szabolcs, P.; Steinberg, H. N.; Bergerson, J.; Skoskiewicz, A.; Makhija, M.; Davis, J.; Foruraghi, L.; Palmer, C.; Fuleihan, R. L.; Church, J. A.; Bhandoola, A.; Lionakis, M. S.; Campbell, S.; Leto, T. L.; Kuhns, D. B.; Holland, S. M. Dominant activating RAC2 mutation with lymphopenia, immunodeficiency, and cytoskeletal defects. Blood 2019, 133, 1977–1988. DOI: 10.1182/blood-2018-11-886028.

(21) Liu, Y.; Cheng, G.; Song, Z.; Xu, T.; Ruan, H.; Cao, Q.; Wang, K.; Bao, L.; Liu, J.; Zhou, L.; Liu, D.; Yang, H.; Chen, K.; Zhang, X. RAC2 acts as a prognostic biomarker and promotes the progression of clear cell renal cell carcinoma. Int J Oncol 2019, 55, 645–656. DOI: 10.3892/ijo.2019.4849.

(22) Navarro-Lerida, I.; Sanchez-Perales, S.; Calvo, M.; Rentero, C.; Zheng, Y.; Enrich, C.; Del Pozo, M. A. A palmitoylation switch mechanism regulates Rac1 function and membrane organization. EMBO J 2012, 31, 534–551. DOI: 10.1038/emboj.2011.446.

(23) Singh, N. K.; Janjanam, J.; Rao, G. N. p115 RhoGEF activates the Rac1 GTPase signaling cascade in MCP1 chemokine-induced vascular smooth muscle cell migration and proliferation. J Biol Chem 2017, 292, 14080–14091. DOI: 10.1074/jbc.M117.777896.

(24) Muller, P. M.; Rademacher, J.; Bagshaw, R. D.; Wortmann, C.; Barth, C.; van Unen, J.; Alp, K. M.; Giudice, G.; Eccles, R. L.; Heinrich, L. E.; Pascual-Vargas, P.; Sanchez-Castro, M.; Brandenburg, L.; Mbamalu, G.; Tucholska, M.; Spatt, L.; Czajkowski, M. T.; Welke, R. W.; Zhang, S.; Nguyen, V.; Rrustemi, T.; Trnka, P.; Freitag, K.; Larsen, B.; Popp, O.; Mertins, P.; Gingras, A. C.; Roth, F. P.; Colwill, K.; Bakal, C.; Pertz, O.; Pawson, T.; Petsalaki, E.; Rocks, O. Systems analysis of RhoGEF and RhoGAP regulatory proteins reveals spatially organized RAC1 signalling from integrin adhesions. Nat Cell Biol 2020, 22, 498–511. DOI: 10.1038/s41556-020-0488-x.

(25) Berta, D.; Gehrke, S.; Nyiri, K.; Vertessy, B. G.; Rosta, E. Mechanism-Based Redesign of GAP to Activate Oncogenic Ras. J Am Chem Soc 2023, 145, 20302–20310. DOI: 10.1021/jacs.3c04330.

(26) Liu, J.; Zhang, J.; Yang, Y.; Huang, H.; Shen, W.; Hu, Q.; Wang, X.; Wu, J.; Shi, Y. Conformational change upon ligand binding and dynamics of the PDZ domain from leukemia-associated Rho guanine nucleotide exchange factor. Protein Sci 2008, 17, 1003–1014. DOI: 10.1110/ps.073416508.

(27) Chichili, V. P. R.; Chew, T. W.; Shankar, S.; Er, S. Y.; Chin, C. F.; Jobichen, C.; Qiurong Pan, C.; Zhou, Y.; Yeong, F. M.; Low, B. C.; Sivaraman, J. Structural basis for p50RhoGAP BCH domain-mediated regulation of Rho inactivation. Proc Natl Acad Sci U S A 2021, 118, e2014242118. DOI: 10.1073/pnas.2014242118.

(28) Xu, L.; Jang, H.; Nussinov, R. Allosteric modulation of NF1 GAP: Differential distributions of catalytically competent populations in loss-of-function and gain-of-function mutants. Protein Sci 2025, 34, e70042. DOI: 10.1002/pro.70042.

(29) Kotting, C.; Kallenbach, A.; Suveyzdis, Y.; Wittinghofer, A.; Gerwert, K. The GAP arginine finger movement into the catalytic site of Ras increases the activation entropy. Proc Natl Acad Sci U S A 2008, 105, 6260–6265. DOI: 10.1073/pnas.0712095105.

(30) Patel, L. A.; Waybright, T. J.; Stephen, A. G.; Neale, C. GAP positions catalytic H-Ras residue Q61 for GTP hydrolysis in molecular dynamics simulations, complicating chemical rescue of Ras deactivation. Comput Biol Chem 2023, 104, 107835. DOI: 10.1016/j.compbiolchem.2023.107835.

(31) Ahmadian, M. R.; Wiesmuller, L.; Lautwein, A.; Bischoff, F. R.; Wittinghofer, A. Structural differences in the minimal catalytic domains of the GTPase-activating proteins p120GAP and neurofibromin. J Biol Chem 1996, 271, 16409–16415. DOI: 10.1074/jbc.271.27.16409.

(32) Doye, A.; Chaintreuil, P.; Lagresle-Peyrou, C.; Batistic, L.; Marion, V.; Munro, P.; Loubatier, C.; Chirara, R.; Sorel, N.; Bessot, B.; Bronnec, P.; Contenti, J.; Courjon, J.; Giordanengo, V.; Jacquel, A.; Barbry, P.; Couralet, M.; Aladjidi, N.; Fischer, A.; Cavazzana, M.; Mallebranche, C.; Visvikis, O.; Kracker, S.; Moshous, D.; Verhoeyen, E.; Boyer, L. RAC2 gain-of-function variants causing inborn error of immunity drive NLRP3 inflammasome activation. J Exp Med 2024, 221, e20231562. DOI: 10.1084/jem.20231562.

(33) Gu, Y.; Jia, B.; Yang, F. C.; D’Souza, M.; Harris, C. E.; Derrow, C. W.; Zheng, Y.; Williams, D. A. Biochemical and biological characterization of a human Rac2 GTPase mutant associated with phagocytic immunodeficiency. J Biol Chem 2001, 276, 15929–15938. DOI: 10.1074/jbc.M010445200.

(34) Lagresle-Peyrou, C.; Olichon, A.; Sadek, H.; Roche, P.; Tardy, C.; Da Silva, C.; Garrigue, A.; Fischer, A.; Moshous, D.; Collette, Y.; Picard, C.; Casanova, J. L.; Andre, I.; Cavazzana, M. A gain-of-function RAC2 mutation is associated with bone-marrow hypoplasia and an autosomal dominant form of severe combined immunodeficiency. Haematologica 2021, 106, 404–411. DOI: 10.3324/haematol.2019.230250.

(35) Alkhairy, O. K.; Rezaei, N.; Graham, R. R.; Abolhassani, H.; Borte, S.; Hultenby, K.; Wu, C.; Aghamohammadi, A.; Williams, D. A.; Behrens, T. W.; Hammarstrom, L.; Pan-Hammarstrom, Q. RAC2 loss-of-function mutation in 2 siblings with characteristics of common variable immunodeficiency. J Allergy Clin Immunol 2015, 135, 1380–1384 e1381–1385. DOI: 10.1016/j.jaci.2014.10.039.

(36) Zhang, L.; Lv, G.; Peng, Y.; Yang, L.; Chen, J.; An, Y.; Zhang, Z.; Tang, X.; Li, Z.; Zhao, X. A Novel RAC2 Mutation Causing Combined Immunodeficiency. J Clin Immunol 2023, 43, 229–240. DOI: 10.1007/s10875-022-01373-8.

(37) Abell, A. N.; DeCathelineau, A. M.; Weed, S. A.; Ambruso, D. R.; Riches, D. W.; Johnson, G. L. Rac2D57N, a dominant inhibitory Rac2 mutant that inhibits p38 kinase signaling and prevents surface ruffling in bone-marrow-derived macrophages. J Cell Sci 2004, 117, 243–255. DOI: 10.1242/jcs.00853.

(38) Donko, A.; Sharapova, S. O.; Kabat, J.; Ganesan, S.; Hauck, F. H.; Bergerson, J. R. E.; Marois, L.; Abbott, J.; Moshous, D.; Williams, K. W.; Campbell, N.; Martin, P. L.; Lagresle-Peyrou, C.; Trojan, T.; Kuzmenko, N. B.; Deordieva, E. A.; Raykina, E. V.; Abers, M. S.; Abolhassani, H.; Barlogis, V.; Milla, C.; Hall, G.; Mousallem, T.; Church, J.; Kapoor, N.; Cros, G.; Chapdelaine, H.; Franco-Jarava, C.; Lopez-Lerma, I.; Miano, M.; Leiding, J. W.; Klein, C.; Stasia, M. J.; Fischer, A.; Hsiao, K. C.; Martelius, T.; Seppanen, M. R. J.; Barmettler, S.; Walter, J.; Masmas, T. N.; Mukhina, A. A.; Falcone, E. L.; Kracker, S.; Shcherbina, A.; Holland, S. M.; Leto, T. L.; Hsu, A. P. Clinical and functional spectrum of RAC2-related immunodeficiency. Blood 2024, 143, 1476–1487. DOI: 10.1182/blood.2023022098.

(39) Arrington, M. E.; Temple, B.; Schaefer, A.; Campbell, S. L. The molecular basis for immune dysregulation by the hyperactivated E62K mutant of the GTPase RAC2. J Biol Chem 2020, 295, 12130–12142. DOI: 10.1074/jbc.RA120.012915.

(40) Mishra, A. K.; Rodriguez, M.; Torres, A. Y.; Smith, M.; Rodriguez, A.; Bond, A.; Morrissey, M. A.; Montell, D. J. Hyperactive Rac stimulates cannibalism of living target cells and enhances CAR-M-mediated cancer cell killing. Proc Natl Acad Sci U S A 2023, 120, e2310221120. DOI: 10.1073/pnas.2310221120.

(41) Smits, B. M.; Lelieveld, P. H. C.; Ververs, F. A.; Turkenburg, M.; de Koning, C.; van Dijk, M.; Leavis, H. L.; Boelens, J. J.; Lindemans, C. A.; Bloem, A. C.; van de Corput, L.; van Montfrans, J.; Nierkens, S.; van Gijn, M. E.; Geerke, D. P.; Waterham, H. R.; Koenderman, L.; Boes, M. A dominant activating RAC2 variant associated with immunodeficiency and pulmonary disease. Clin Immunol 2020, 212, 108248. DOI: 10.1016/j.clim.2019.108248.

(42) Jang, H.; Yavuz, B. R.; Zhang, M.; Liu, Y.; Nussinov, R. Oncogenic PI3Kalpha variants reveal graded conformational spectrum with mutation-specific cryptic pockets. Commun Chem 2026, 9. DOI: 10.1038/s42004-026-01906-x.

(43) Xu, L.; Eren, M.; Weako, J.; Jang, H.; Keskin, O.; Gursoy, A.; Nussinov, R. The structural heterogeneity of AKT autoinhibition. Protein Sci 2026, 35, e70420. DOI: 10.1002/pro.70420.

(44) Xu, L.; Liu, Y.; Jang, H.; Nussinov, R. M-Ras distinct activation scenarios: A mechanistic outlook and targeting. Comput Struct Biotechnol J 2025, 27, 5207–5219. DOI: 10.1016/j.csbj.2025.11.025.

(45) Haspel, N.; Jang, H.; Nussinov, R. Allosteric Activation of RhoA Complexed with p115-RhoGEF Deciphered by Conformational Dynamics. J Chem Inf Model 2024, 64, 862–873. DOI: 10.1021/acs.jcim.3c01412.

(46) Jang, H.; Chen, J.; Iakoucheva, L. M.; Nussinov, R. Cancer and Autism: How PTEN Mutations Degrade Function at the Membrane and Isoform Expression in the Human Brain. J Mol Biol 2023, 435, 168354. DOI: 10.1016/j.jmb.2023.168354.

(47) Haspel, N.; Jang, H.; Nussinov, R. Active and Inactive Cdc42 Differ in Their Insert Region Conformational Dynamics. Biophys J 2021, 120, 306–318. DOI: 10.1016/j.bpj.2020.12.007.

(48) Brooks, B. R.; Brooks, C. L., 3rd; Mackerell, A. D., Jr.; Nilsson, L.; Petrella, R. J.; Roux, B.; Won, Y.; Archontis, G.; Bartels, C.; Boresch, S.; Caflisch, A.; Caves, L.; Cui, Q.; Dinner, A. R.; Feig, M.; Fischer, S.; Gao, J.; Hodoscek, M.; Im, W.; Kuczera, K.; Lazaridis, T.; Ma, J.; Ovchinnikov, V.; Paci, E.; Pastor, R. W.; Post, C. B.; Pu, J. Z.; Schaefer, M.; Tidor, B.; Venable, R. M.; Woodcock, H. L.; Wu, X.; Yang, W.; York, D. M.; Karplus, M. CHARMM: the biomolecular simulation program. J Comput Chem 2009, 30, 1545–1614. DOI: 10.1002/jcc.21287.

(49) Huang, J.; Rauscher, S.; Nawrocki, G.; Ran, T.; Feig, M.; de Groot, B. L.; Grubmuller, H.; MacKerell, A. D., Jr. CHARMM36m: an improved force field for folded and intrinsically disordered proteins. Nat Methods 2017, 14, 71–73. DOI: 10.1038/nmeth.4067.

(50) Klauda, J. B.; Venable, R. M.; Freites, J. A.; O’Connor, J. W.; Tobias, D. J.; Mondragon-Ramirez, C.; Vorobyov, I.; MacKerell, A. D., Jr.; Pastor, R. W. Update of the CHARMM all-atom additive force field for lipids: validation on six lipid types. J Phys Chem B 2010, 114, 7830–7843. DOI: 10.1021/jp101759q.

(51) Phillips, J. C.; Braun, R.; Wang, W.; Gumbart, J.; Tajkhorshid, E.; Villa, E.; Chipot, C.; Skeel, R. D.; Kale, L.; Schulten, K. Scalable molecular dynamics with NAMD. J Comput Chem 2005, 26, 1781–1802. DOI: 10.1002/jcc.20289.

(52) Grudzien, P.; Jang, H.; Leschinsky, N.; Nussinov, R.; Gaponenko, V. Conformational Dynamics Allows Sampling of an "Active-like" State by Oncogenic K-Ras-GDP. J Mol Biol 2022, 434, 167695. DOI: 10.1016/j.jmb.2022.167695.

(53) Rudack, T.; Xia, F.; Schlitter, J.; Kotting, C.; Gerwert, K. Ras and GTPase-activating protein (GAP) drive GTP into a precatalytic state as revealed by combining FTIR and biomolecular simulations. Proc Natl Acad Sci U S A 2012, 109, 15295–15300. DOI: 10.1073/pnas.1204333109.

(54) Calixto, A. R.; Moreira, C.; Pabis, A.; Kotting, C.; Gerwert, K.; Rudack, T.; Kamerlin, S. C. L. GTP Hydrolysis Without an Active Site Base: A Unifying Mechanism for Ras and Related GTPases. J Am Chem Soc 2019, 141, 10684–10701. DOI: 10.1021/jacs.9b03193.

(55) Nussinov, R.; Regev, C.; Jang, H. Energy landscapes in molecular biology: History, principles, and perspectives. Q Rev Biophys 2026, 59, e7. DOI: 10.1017/S0033583526100134.

(56) Nussinov, R.; Regev, C.; Jang, H. Leveraging conformational ensembles in allosteric drug discovery. Trends Pharmacol Sci 2026, 47, 276–289. DOI: 10.1016/j.tips.2026.01.006.

(57) Nussinov, R.; Yavuz, B. R.; Jang, H. Allostery: allosteric networks and allosteric signaling bias. Q Rev Biophys 2025, 58, e17. DOI: 10.1017/S0033583525100061.

(58) Brown, K. M.; Xu, M.; Sargen, M.; Jang, H.; Zhang, M.; Zhang, T.; Zhu, B.; Jones, K.; Kim, J.; Mendoza, L.; Hayward, N. K.; Tucker, M. A.; Goldstein, A. M.; Yang, X. R.; Stewart, D. R.; Hicks, B.; Consonni, D.; Pesatori, A. C.; Fargnoli, M. C.; Peris, K.; Stratigos, A.; Menin, C.; Ghiorzo, P.; Puig, S.; Nagore, E.; MelaNostrum, C.; Andresson, T.; Nussinov, R.; Calista, D.; Landi, M. T. Novel MAPK/AKT-impairing germline NRAS variant identified in a melanoma-prone family. Fam Cancer 2022, 21, 347–355. DOI: 10.1007/s10689-021-00267-9.

(59) He, J.; Liu, X.; Zhu, C.; Zha, J.; Li, Q.; Zhao, M.; Wei, J.; Li, M.; Wu, C.; Wang, J.; Jiao, Y.; Ning, S.; Zhou, J.; Hong, Y.; Liu, Y.; He, H.; Zhang, M.; Chen, F.; Li, Y.; He, X.; Wu, J.; Lu, S.; Song, K.; Lu, X.; Zhang, J. ASD2023: towards the integrating landscapes of allosteric knowledgebase. Nucleic Acids Res 2024, 52, D376–D383. DOI: 10.1093/nar/gkad915.

(60) Li, M.; Lan, X.; Shi, X.; Zhu, C.; Lu, X.; Pu, J.; Lu, S.; Zhang, J. Delineating the stepwise millisecond allosteric activation mechanism of the class C GPCR dimer mGlu5. Nat Commun 2024, 15, 7519. DOI: 10.1038/s41467-024-51999-y.

