## Supplemental Figures S1-S7 for "How Functional Variants Reconfigure the Rac2 Conformational Landscape"

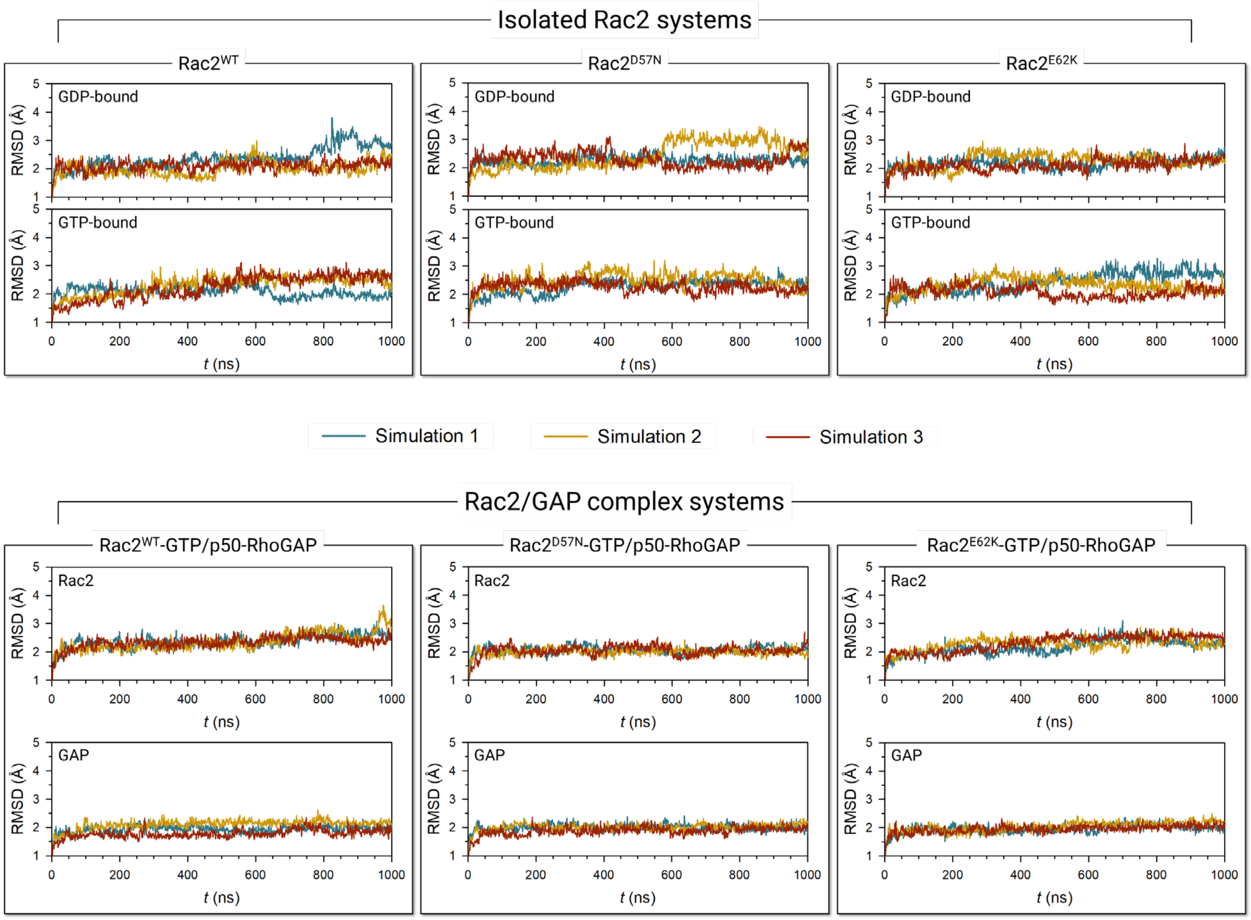


**Figure S1.** Time series showing the root-mean-square-deviations (RMSDs) for three replica simulations of the isolated Rac2$^{WT}$, Rac2$^{D57N}$, and Rac2$^{E62K}$ in the GDP- and GTP-bound states (*top row*). The same for three replica simulations of the GTP-bound Rac2$^{WT}$, Rac2$^{D57N}$, and Rac2$^{E62K}$ in complex with p50-RhoGAP (*bottom row*).

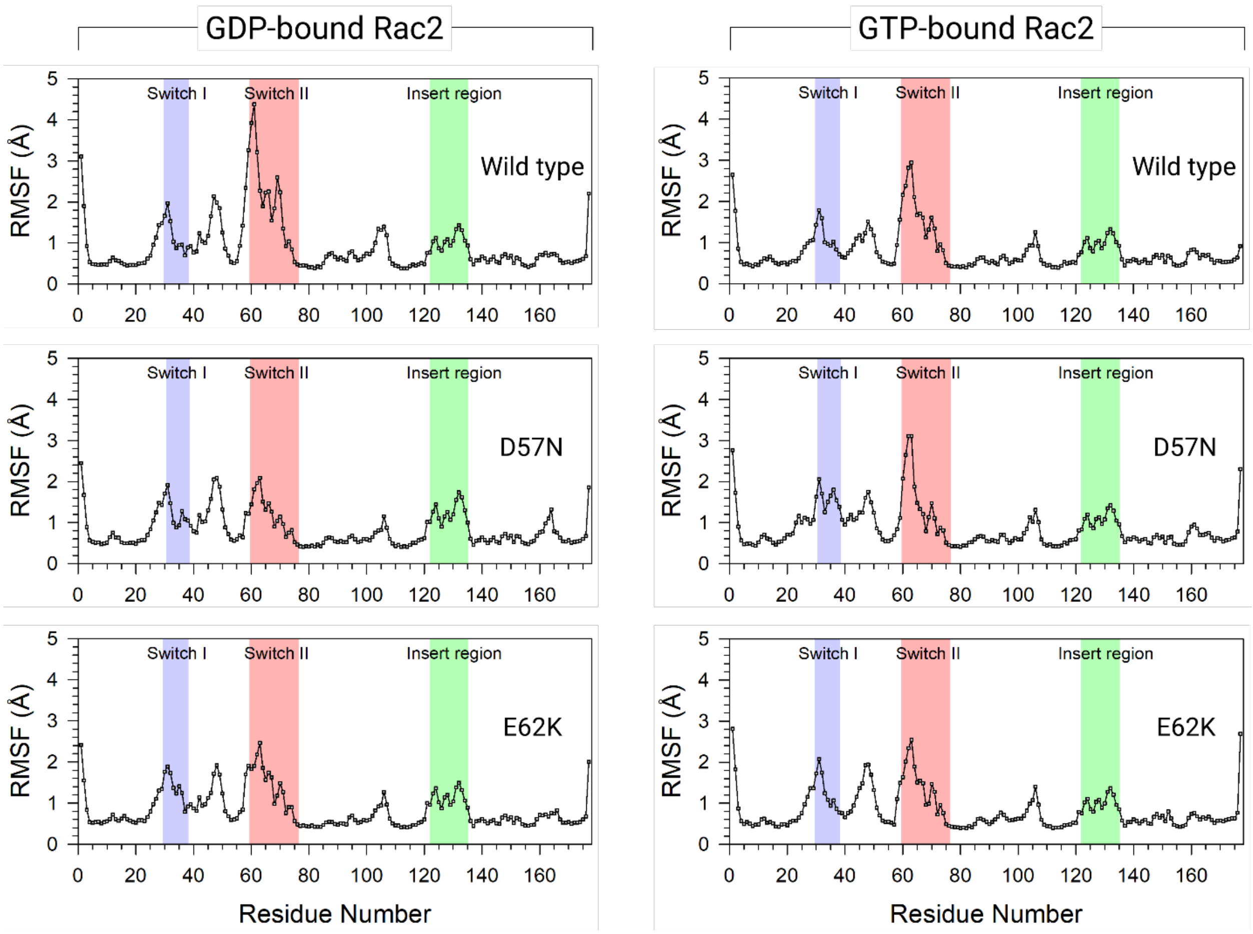


**Figure S2.** The root mean-square fluctuations (RMSFs) of Rac2$^{WT}$, Rac2$^{D57N}$, and Rac2$^{E62K}$ for the GDP-bound (*left column*) and the GTP-bound (*right column*) states as a function of the residue number. Light blue, light red, and light green backgrounds denote the Switch I, Switch II, and the insert regions, respectively.

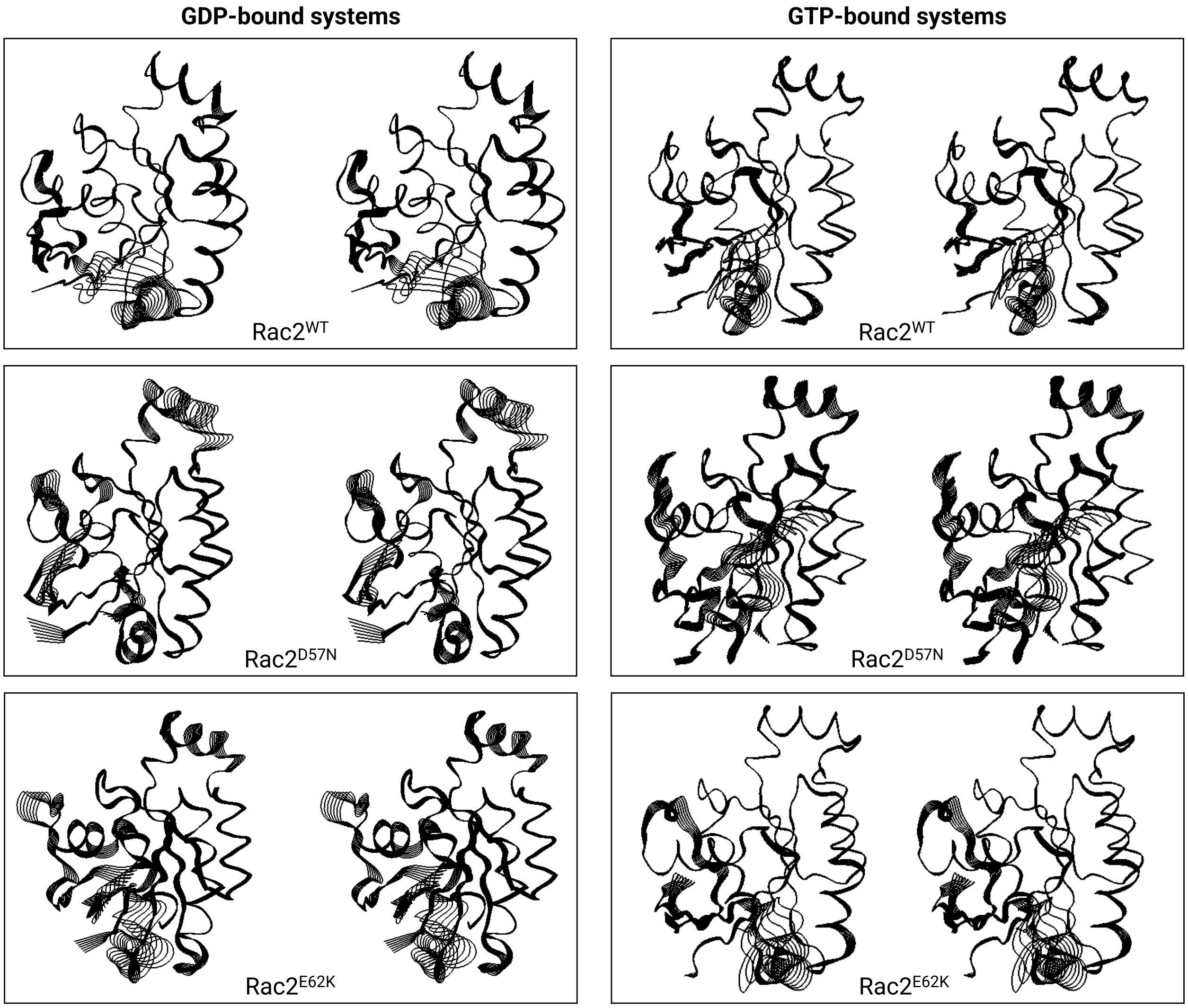


**Figure S3.** Superimpositions of protein motion at the main normal mode with the lowest-frequency motion in a stereo pair view for Rac2$^{WT}$, Rac2$^{D57N}$, and Rac2$^{E62K}$ in the GDP-bound (*left column*) and the GTP-bound (*right column*) states. The images show the full range of motion for each residue. The motion clearly demonstrates the conformational flexibility of both switch regions in all systems. The GDP- and GTP-bound D57N mutants exhibit significant motion in the insert region compared to the other systems.

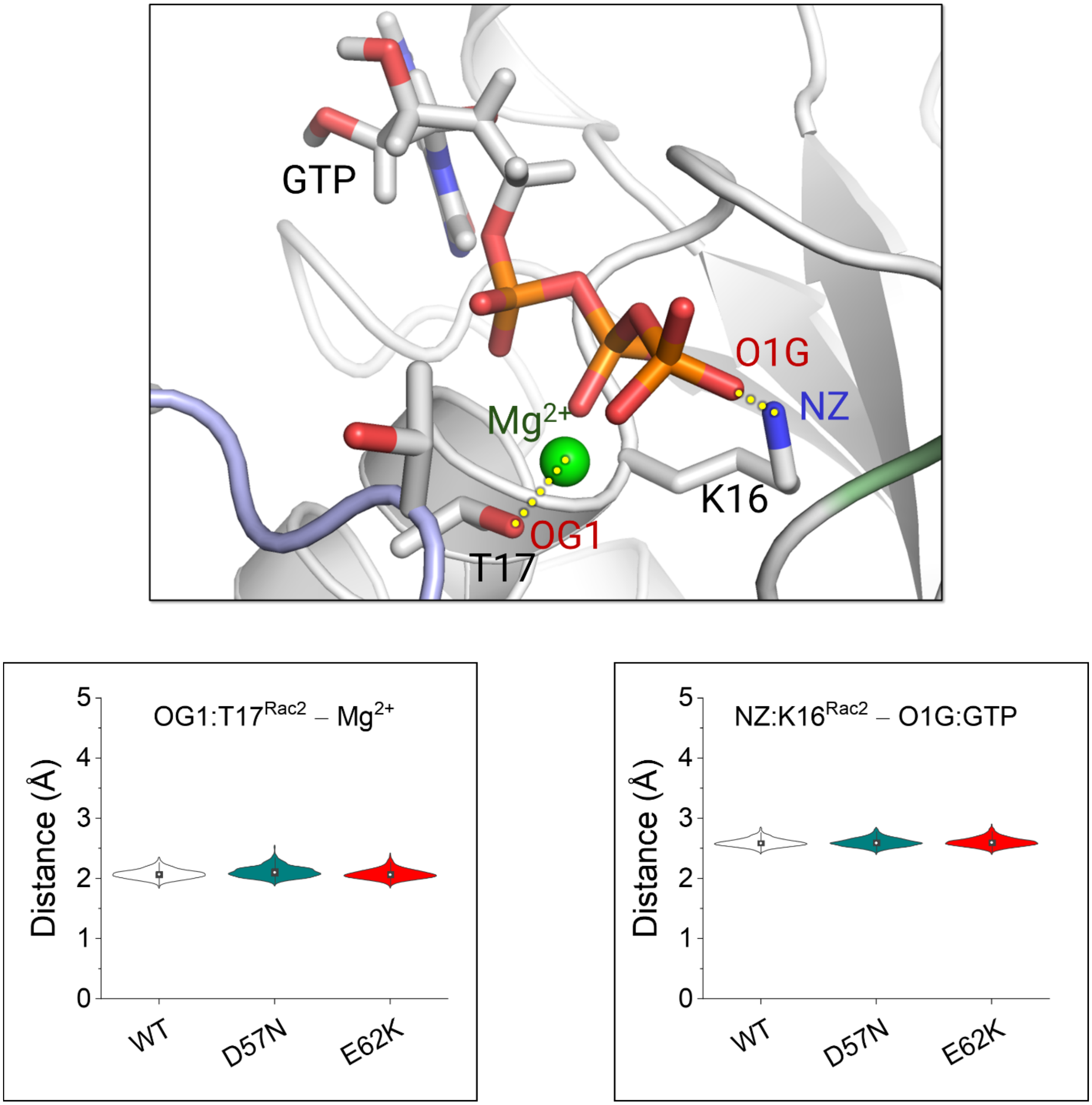


**Figure S4.** An example of snapshot highlighting the Thr17–$Mg^{2+}$ coordination and the Lys16–GTP salt bridge in the isolated $Rac2^{WT}$-GTP (*top panel*). The violin plots show the atomic pair distances between Thr17-OG1 and $Mg^{2+}$ (*bottom left*), and between Lys16-NZ and GTP-O1G (*bottom right*). These distances are highly conserved.

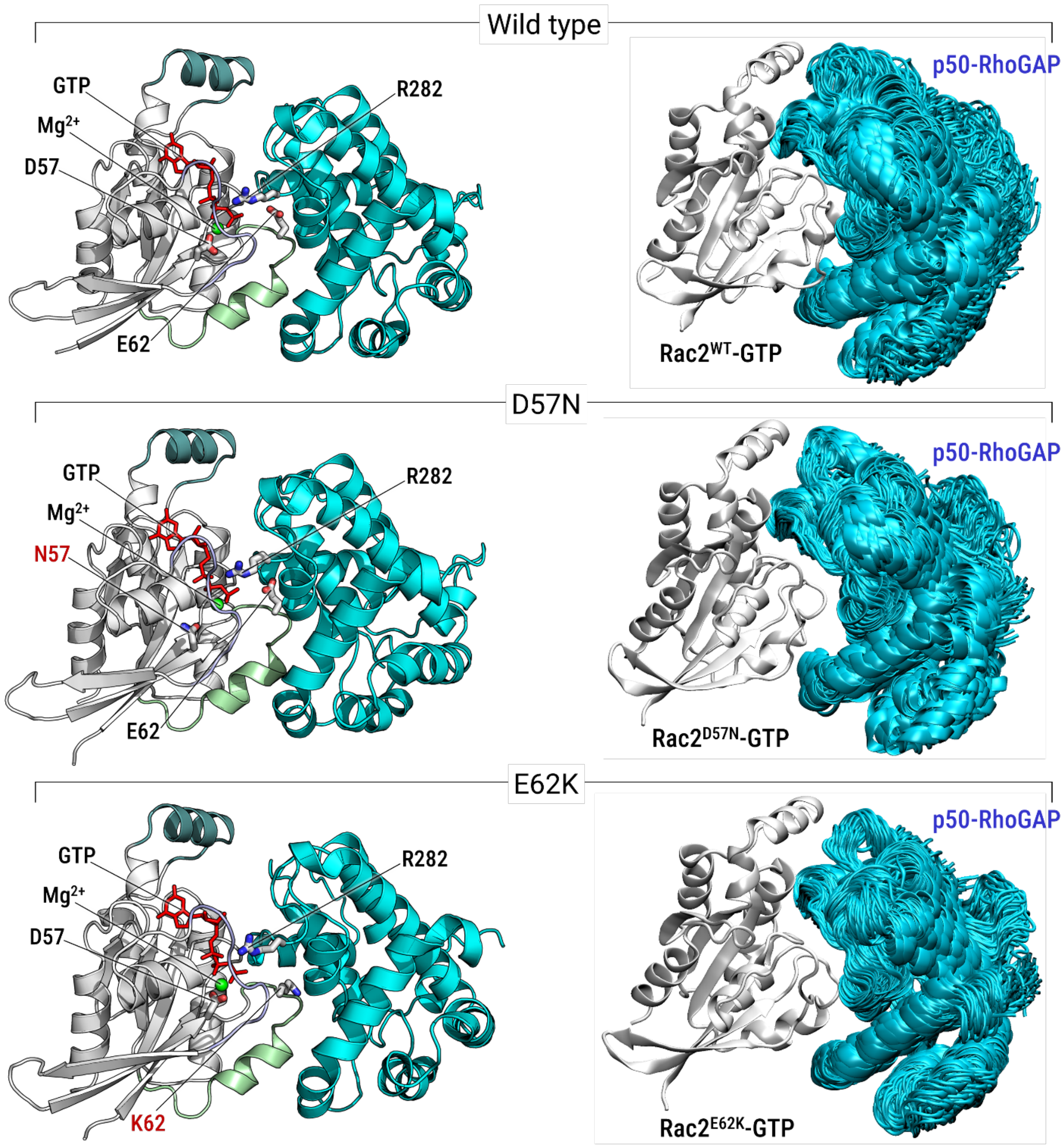


**Figure S5**. Snapshots depicting the initial conformations of the Rac2/p50-RhoGAP complex in cartoon representation, and the structural alignment of simulated p50-RhoGAP with respect to GTP-bound wild-type Rac2 and its D57N and E62K mutants. The sticks highlight the D57N and E62K mutant residues on Rac2, as well as the Arg282 finger on the GAP.

**NF1 GRD (isoform2)**

1210 1220 1230 1240 1250 1260 1270
....FERLVE LVTMMGDQGE LPIAMALANV VPCSQWDELA RVLVTLFDSR HLLYQLLWNM FSKEVELADS
1280 1290 1300 1310 1320 1330 1340
MQTLFRGNSL ASKIMTFCFK VYGATYLQKL LDPLLRIVIT SSDWQHVSFE VDPTRLEPSE SLEENQRNLL
1350 1360 1370 1380 1390 1400 1410
QMTEKFFHAI ISSSSEFPPQ LRSVCHCLYQ VVSQRFPQNS IGAVGSAMFL RFINPAIVSP YEAGILDKKP
1420 1430 1440 1450 1460 1470 1480
PPRIERGLKL MSKILQSIAN HVLFTKEEHM RPFNDFVKSN FDAARRFFLD IASDCPTSDA VNHSLSFISD
1490 1500 1510 1520 1530
GNVLALHRLL WNNQEKIGQY LSSNRDHKAV GRRPFDKMAT LLAYLGPPEH

**p50-RhoGAP**

240 250 260 270 280 290 300
......LPNQ QFGVSLQHLQ EKNPEQEPIP IVLRETVAYL QAHALTTEGI FRRSANTQVV REVQQKYNMG
310 320 330 340 350 360 370
LPVDFDQYNE LHLPAVILKT FLRELPEPLL TFDLYPHVVG FLNIDESQRV PATLQVLQTL PEENYQVLRF
380 390 400 410 420 430
LTAFLVQISA HSDQNKMTNT NLAVVFGPNL LWAKDAAITL KAINPINTFT KFLLDHQGEL FPS......

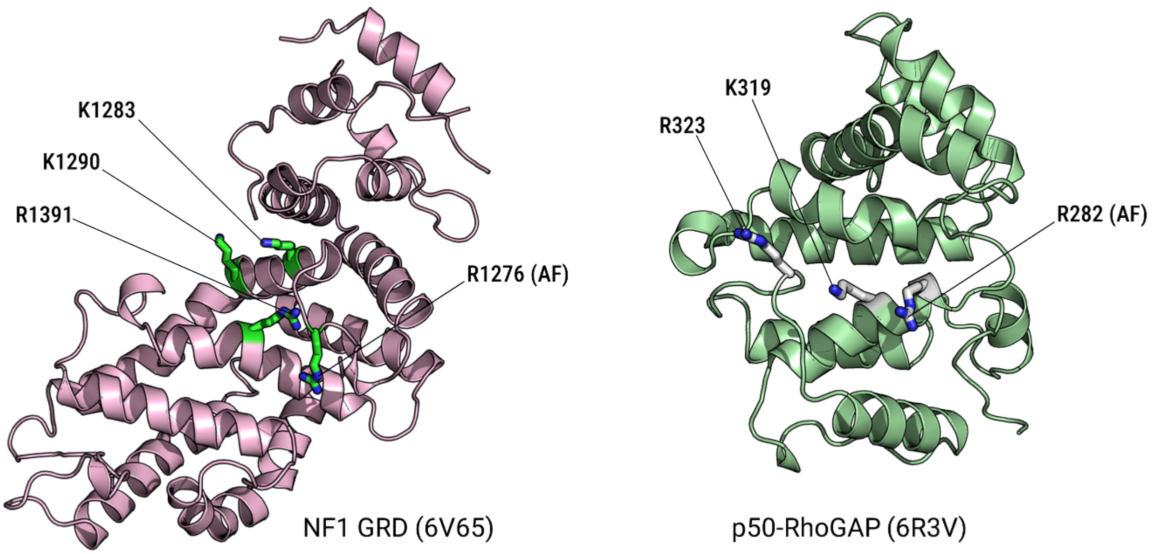


**Figure S6.** Sequence comparison between the NF1 GTPase-activating protein-related domain (GRD, isoform2) and p50-RhoGAP. In the sequences, the key basic residues at the interface with GTPases are colored blue. The cartoons depict the crystal structures of the NF1 GRD (PDB ID: 6V65) and the p50-RhoGAP (PDB ID: 6R3V), highlighting the key basic residues and the arginine finger (AF) as sticks.

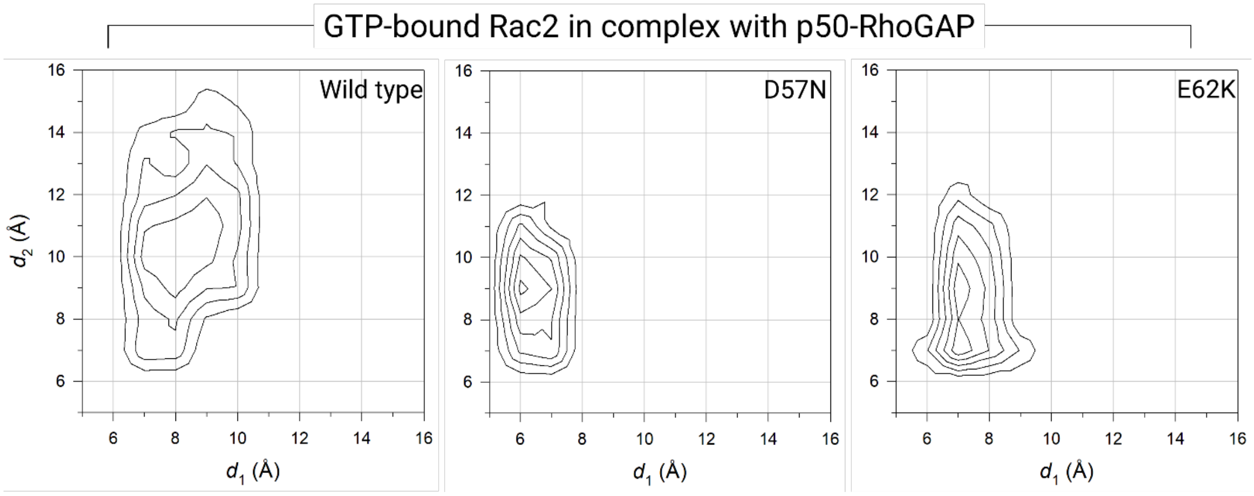


**Figure S7.** The two-dimensional potential of mean force, $\Delta G_{PMF}$, representing the relative free energy profile along reaction coordinates, $d_1$ and $d_2$. In the calculation of the probability distributions for two atom pair distances, $d_1$ is defined as the distance from Gly60-Cα to GTP-$P_\beta$ and $d_2$ is defined as the distance from Thr35-Cα to GTP-$P_\beta$. The contour plots illustrate the distance profiles for the GTP-bound $Rac2^{WT}$, $Rac2^{D57N}$, and $Rac2^{E62K}$ in complex with p50-RhoGAP.
